# CRISPR interference functional genomics of coding and non-coding determinants of *Bacillus subtilis* biofilms

**DOI:** 10.64898/2026.06.23.734000

**Authors:** Hadrien H. Barras, Pierre Nicolas, Despoina Anastasopoulou, Romain Briandet, Marie-Françoise Noirot-Gros

**Affiliations:** Université Paris-Saclay, INRAE, AgroParisTech, Micalis Institute, 78350 Jouy-en-Josas, France; Université Paris-Saclay, INRAE, MaIAGE, 78350 Jouy-en-Josas, France

## Abstract

The architecture of *Bacillus subtilis* biofilms is influenced by the coordinated regulation of cellular specialization, matrix assembly, and metabolism. *B. subtilis* can form different types of biofilm in diverse physical and chemical environments. Understanding the molecular mechanisms that drive biofilm heterogeneity and adaptation to different environmental niches is crucial for developing more effective strategies to control their formation. In this study, we developed a tightly dual-regulated CRISPR interference (CRISPRi) system and employed multi-scale imaging to investigate the functions of individual genes in two distinct biofilm models: the floating pellicle and the intricate, three-dimensionally structured macrocolony, which develop at the liquid-air and solid-air interfaces, respectively. Our findings validated the CRISPRi approach as a powerful method for studying biofilm development over extended periods and revealed that numerous small non-coding RNAs are involved in regulating biofilm growth dynamics and architecture. The CRISPRi approach was also applied to a pool of 507 genes and transcription units, including protein-coding genes and non-coding RNAs, to screen for cell fitness in these two biofilm models. We discovered that, while both biofilm forms rely on fundamental processes such as cell wall synthesis and nucleotide metabolism, they exhibit different genetic dependencies with regard to matrix composition, motility, and signaling. Exopolysaccharide production, motility, and chemotaxis are crucial for pellicle formation. In contrast, macrocolony development is influenced by γ-polyglutamate synthesis and nutrient acquisition functions. Genes of unknown function were also identified to play a differentially important role in the two biofilm forms. Additionally, the CRISPRi screens revealed further non-coding RNAs regulating biofilm architecture and growth dynamics, adding to the existing layers of post-transcriptional control. Collectively, these results demonstrate that biofilm formation at different physical interfaces is governed by a combination of shared and unique genetic pathways tailored to the specific biofilm environment, thereby opening research avenues into the molecular mechanisms specific to the solid-air and liquid-air interfaces.

## Introduction

Bacterial biofilms are heterogeneous assemblages of cells exhibiting various morphologies and traits that facilitate their adaptation and survival in fluctuating environments ^1^. Despite significant progress in our understanding of biofilm biology, the genetic components and molecular pathways responsible for shaping the different types of biofilms formed at various interfaces remain elusive. This knowledge gap hinders our understanding of the mechanisms that govern biofilm diversity. *Bacillus subtilis* is a well-established Gram-positive model organism that has been extensively studied for its capacity to form biofilms ^2-4^. *B. subtilis* biofilms develop in distinct morphotypes, notably as macrocolonies at solid-air interfaces and as pellicles at liquid-air interfaces. Each type of biofilm has unique architectural features and gene expression profiles that reflect specialized physiological adaptations optimized for survival and resource utilization within its respective ecological niche ^3^.

The architecture of *B. subtilis* biofilms primarily depends on the coordinated production of extracellular matrix components. Key constituents of the matrix include exopolysaccharides (EPS), encoded by the large *epsA-O* operon, which serve as the primary scaffold of the biofilm matrix ^4-6^. Amyloid fibers, primarily composed of the TasA protein, provide mechanical strength. The production of amyloid fibers by TasA is dependent on all three elements of the *tapA-sipW-tasA* operon ^5^. TasA and its chaperone accessory protein TapA are secreted into the matrix by the membrane-bound signal peptidase SipW, where TasA assembles into fibers ^7-9^. Concurrently, hydrophobins such as BslA enhance surface hydrophobicity. BslA is a secreted protein with self-assembly properties that forms a highly ordered protective “raincoat” around the biofilm matrix ^10-12^. Together, these components provide structural integrity, cohesion, and protection ^3-5,10,11,13^. Their expression and assembly are regulated at multiple genetic and post-transcriptional levels, ensuring the dynamic modulation of biofilm architecture in response to environmental cues.

The transition of *B. subtilis* from a planktonic-motile state to a biofilm-sessile state is initiated by a switch orchestrated by a multifaceted signaling pathway involving the global transition regulators Spo0A and AbrB and dedicated biofilm regulators ^14-18^. Biofilm formation depends on the level of phosphorylated Spo0A, which is regulated by a network of histidine kinases (KinA-KinE) and phosphatases ^19-22^. These components trigger a phosphorelay cascade in response to various environmental signals, thereby maintaining moderate levels of Spo0A-P in cells, which, in turn, regulates the expression of the genes required for this lifestyle transition ^17,23^. Additional post-translational modifications of proteins by bacterial tyrosine kinases (BY-kinases) have also been reported ^24-26^. The BY-activator/kinase complexes EpsA/EpsB and TkmA/PtkA regulate exopolysaccharide biosynthesis ^27^. Additionally, intracellular mRNA levels are regulated by the endoribonuclease RNase Y, which plays a role in biofilm formation in *B. subtilis* ^28^. Bacterial subpopulations within biofilms adapt and respond uniquely to local chemical environmental conditions, such as nutrient gradients, oxygen levels, the presence of waste products, and signaling compounds. This results in significant physiological heterogeneity across both spatial and temporal scales. Transcriptomic analyses of biofilms have revealed numerous differentially expressed genes ^29-31^. However, whether these genes directly contribute to biofilm development and morphology remains unclear.

In recent years, non-coding RNAs (ncRNAs) have emerged as key regulatory elements in bacterial physiology, including biofilm development ^32^. Although early studies identified only a limited number of ncRNAs involved in *B. subtilis* biofilm formation, transcriptomic analyses have revealed a broad repertoire of ncRNAs that are upregulated during the initial stages of biofilm development ^31,33^. These ncRNAs are distributed across diverse genomic contexts, including untranslated regions, intergenic sequences, and antisense to coding sequences. Their regulatory roles are primarily involve post-transcriptional modulation of gene expression, affecting mRNA stability, translation efficiency, and feedback mechanisms within transcriptional networks ^34^. However, the role of these ncRNAs in the context of *B. subtilis* biofilm formation remains to be elucidated.

*Bacillus subtilis* NDmed is an excellent model for studying biofilms due to its genetic tractability and robust biofilm development ^35^. NDmed can develop a highly intricate, wrinkled macrocolony biofilm on the surface of an agar plate and a thick, structured floating biofilm pellicle at the liquid-air interface. In rich media, *B. subtilis* NDmed can also form submerged biofilms at solid-liquid interfaces, with biphasic dynamics coordinated with pellicle formation ^36^. Its well-characterized genetic background facilitates detailed functional analyses, making it an ideal candidate for dissecting the molecular determinants of biofilm heterogeneity. This study aims to identify genes and non-coding regulatory RNAs that play a significant role in the formation of two distinct biofilm morphotypes: the macrocolony and the floating pellicle. To accomplish this, we employed a CRISPR interference (CRISPRi) approach to specifically repress gene expression ^37^. We implemented a dual regulatory system that combines a transcriptional repressor and a translational riboswitch to tightly control dCas9 expression. This system provided a streamlined approach for studying the functional roles of coding and non-coding genes in biofilm formation. Based on previous transcriptomic data ^36^, we constructed a library of guide RNAs targeting genetic elements upregulated during the initial phase of biofilm formation and performed CRISPRi screens in the pellicle and macrocolony biofilm models. Through these screens, we identified key pathways, including cell wall synthesis, nucleotide metabolism, motility, and extracellular matrix production, with some genes exhibiting biofilm-model specificity. For instance, we found that exopolysaccharide biosynthesis genes played a significant role in pellicle formation, whereas γ-polyglutamate synthesis was more important for the formation of the macrocolony. Motility and chemotaxis genes were important for pellicle formation but were not specifically required for the macrocolony model. Furthermore, ncRNAs identified by CRISPRi may contribute to the fine-tuning of biofilm formation in both models, highlighting the complexity of genetic regulation that extends beyond protein-coding genes.

Overall, combining CRISPRi-mediated gene silencing with real-time phenotypic analysis provided a powerful framework for unraveling the genetic determinants of biofilm architecture and dynamics in *B. subtilis*. This dual-model genetic analysis has advanced our understanding of biofilm heterogeneity and offers potential targets for modulating biofilm formation in various applications.

## Results

### 1. A dual regulatory system for the conditional expression of *dCas9* in *B. subtilis*

To ensure precise regulation of *dcas9* expression, we built a dual-input system that combined a previously characterized cumate-inducible toggle switch, which controls gene expression at the transcriptional level ^38^, with a theophylline-responsive riboswitch, which regulates gene expression at the translational level (Figure 1a) ^38-40^. Transcriptional repression by *cymR* is alleviated in the presence of cumate, which binds to CymR and inhibits its ability to bind DNA at the CuO operator site ^40^. The theophylline riboswitch has been engineered to respond positively to the presence of theophylline ^39^. When theophylline binds to the riboswitch, it triggers a conformational change that exposes the ribosome binding site (RBS), facilitating the recruitment of the translation machinery (Figure 1b). Both components have been shown to function independently in *B. subtilis* ^41,42^.

**Figure 1.**
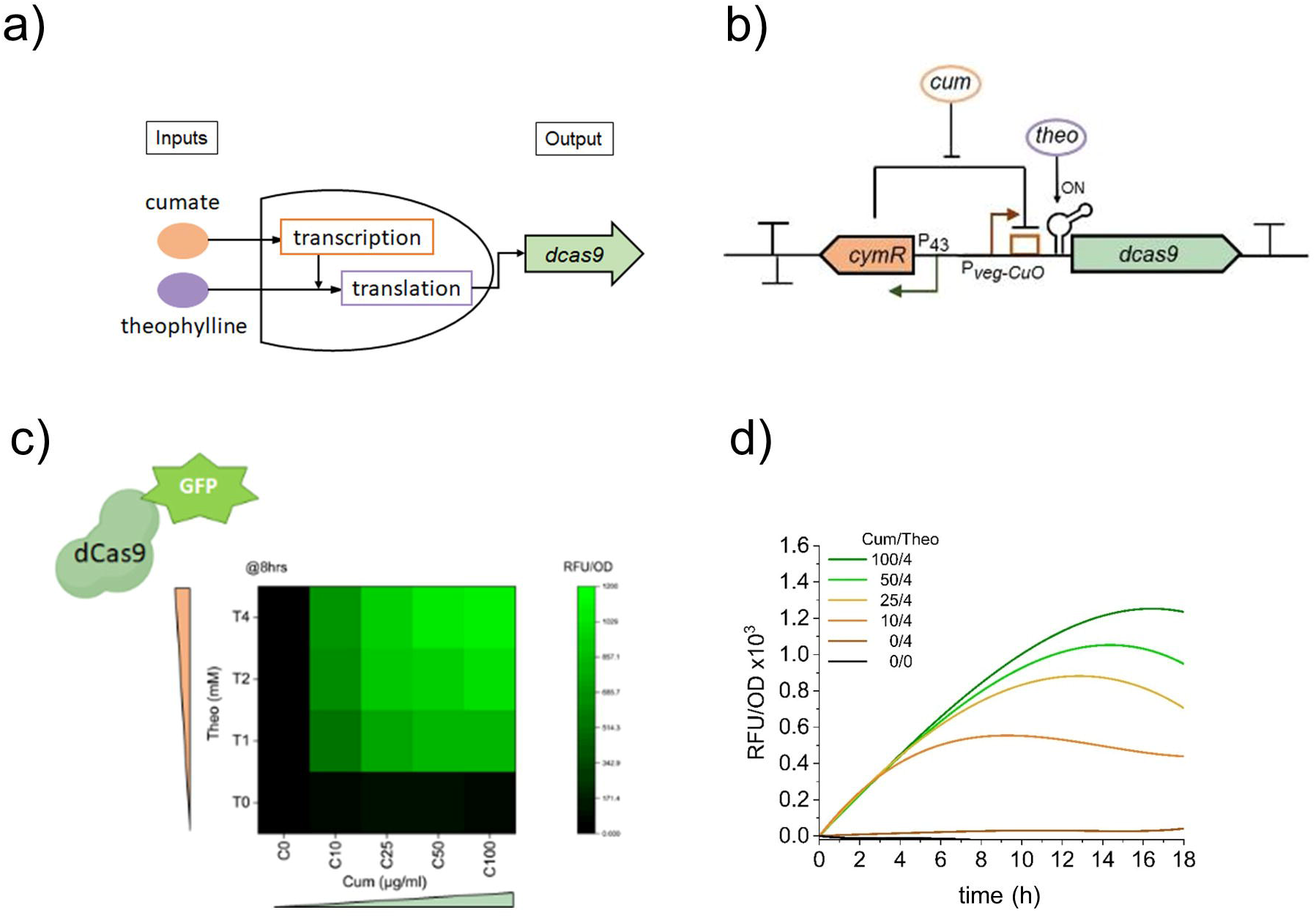
A dual regulatory system for the tight control of expression of dcas9 in *B. subtilis*. (a) Schematic representation of the dual-induced system. (b) Schematic representation of the cumate-theophylline riboswitch dual-control system. The *cymR* gene is expressed under the control of the P43 promoter of *B. subtilis*. In the absence of the chemical compound cumate, the repressor protein CymR binds to its cognate CuO box within the Pveg promoter, preventing transcription. This repression is relieved when cumate binds to CymR, preventing its attachment to the CuO-box. Upon theophylline binding, the riboswitch adopts an ON configuration, releasing the RBS and leading to dCas9 expression. (c-d) Dynamics of dCas9-GFP fusion protein expression in the presence of inducers. NDM101 strain was cultured in the presence of various concentrations of cumate and theophylline. Fluorescence intensities were monitored 8 h after induction by flow cytometry (c) or continuously in a multimodal plate reader (d).

We constructed an NDmed strain (NDM100) carrying a module expressing the *dcas9* gene, which is dually controlled by *cymR* and a theophylline-responsive riboswitch, and stably integrated within the genome at the *lacA* locus. The cumate and theophylline-inducible expression system (CUTE) was evaluated in *B. subtilis* NDmed to assess its response dynamics in the presence or absence of one or both inducers. To achieve this, a translational fusion of the *dCas9* and GFP coding sequences linked by a spacer encoding a glycine-rich sequence was constructed, giving rise to the strain NDM101. In the absence of both inducers, no expression of *dcas9-*GFP above the background was detected, indicating tight repression (Figure 1c). Using a multimodal plate reader, the analysis of green fluorescence levels over time demonstrated a large dynamic range of d*cas9-gfp* expression across various cumate and theophylline concentrations (Figure 1c-d). Flow cytometry revealed that the *dCas9*-GFP protein was expressed homogeneously in the NDM101 population following induction with cumate and theophylline (Figure S1a-b). In the absence of one or both inducers, the fluorescence signal was identical to that of the non-fluorescent NDM100 strain expressing an untagged dCas9 (Figure S1c). These results highlighted the stringency of *dcas9* repression. In the presence of both inducers, fluorescence was detected after one hour and was responsive to increasing concentrations of cumate in the presence of theophylline (Figure S1d).

We then assessed *dCas9* repression efficacy using the CUTE system by monitoring the downregulation of fluorescent reporter genes. For this purpose, we examined the repression of an *mCherry* or a *gfp* gene inserted at the chromosomal *amyE* locus within the NDM100 strain. A guide RNA (g_*mCherry*) was designed to target DNA sequences within the hyperspank promoter for *mCherry* expression, whereas another (g_*gfp*) targeted the beginning of the *gfp* open reading frame (ORF) (Figure S1e-f). We observed that the expression of mCherry remained completely suppressed for a period of 12 h in the presence of both inducers in growing cultures, indicating stable repression over time (Figure S1e). Upon the introduction of inducers into a mid-exponential-phase culture expressing GFP protein (OD = 0.5), a gradual and significant reduction in green fluorescence intensity, attributable to the silencing of the *gfp* gene, was observed by flow cytometry over 5 h (Figure S1f). Collectively, these results demonstrated the effectiveness of the CUTE-dCas9 system in rapidly and efficiently downregulating gene expression over several hours while maintaining tight repression over time.

### 2. CRISPRi-mediated investigation of biofilm formation at solid and liquid-air interfaces

*B. subtilis* NDmed forms highly structured macrocolony biofilms at solid-air on agar surfaces and thick pellicle biofilms at liquid-air interfaces. To evaluate the effectiveness of gene silencing by CRISPRi for studying biofilm formation, we used the CUTE-dCas9 system to observe biofilm phenotypes after silencing genes encoding the main components of the biofilm matrix. Genes from the *eps* operon, as well as *bslA, tapA*, and *tasA*, were knocked down using guide RNAs that targeted either the start of the ORFs, the 5’ untranslated region (5’ UTR), or intergenic sequences (Figure 2). The effects on macrocolony and pellicle phenotypes were monitored after 36 h. In addition, their formation dynamics were observed over 48 or 60 h using automated, real-time, AI-assisted imaging and analysis technology from Reshape Biotech. We first compared the effects of gene knockdown (KD) with those of their gene knockout (KO) counterparts on biofilm phenotypes.

**Figure 2.**
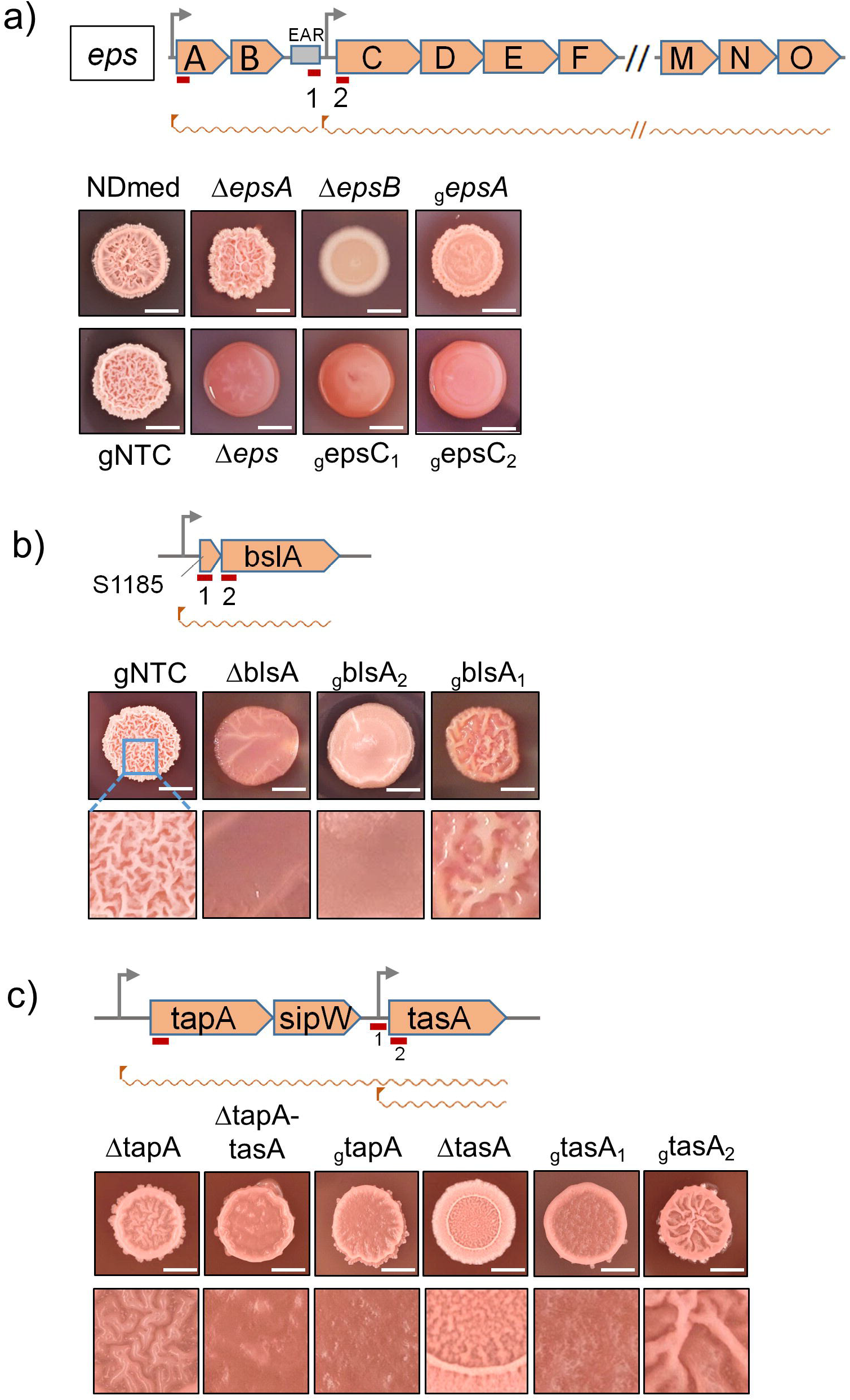
Macrocolony phenotyping of biofilm genes in NDmed. CRISPRi gene knockdowns (KD) were compared to their cognate gene knockouts (KO). (a) Comparative KD and KO of genes from the *eps* operon. The eps operon is pictured, with the target location of the gRNA indicated by red bars. (b) Comparative KD and KO of the *blsA* gene; (c) Comparative KD and KO of the tapA-sigW-tasA operon. The extent of the deleted sequences in the *tasA* gene is indicated. (Scale bar: 5 mm; inside magnification x3)

#### 2.1. Macrocolony formation and morphology

In *B. subtilis*, the 16-kb-long *epsA-O* operon is expressed as two transcription units (TUs) ^31^. The first unit covers the genes *epsA* and *epsB*, which encode a regulatory system composed of a transmembrane modulator and a tyrosine kinase (U2701). The second unit extends from *epsC* to *epsO* (U2699). Complete transcription of this region is facilitated by the EAR regulatory motif located between *epsB* and *epsC* ^43^. Given that the CRISPRi approach operates by downregulating gene expression at the operon level, we silenced these two transcriptional units using RNA guides that target the *epsA* gene at the start of the ORF (g_epsA) and *epsC*, targeted either at the start of the ORF (g_epsC2) or the 5’-proximal of the promoter region, within the EAR region (g_epsC1). After 36 h, macrocolonies with highly wrinkled structures characteristic of NDmed were observed for the NDM100 strain carrying the CUTE-*dCas9* system but expressing a control gRNA that did not target any gene (g_NTC). In contrast, *epsA* knockdown resulted in the formation of a macrocolony devoid of wrinkled structures. This phenotype was in contrast to that of the Δ*epsA* strain, which retained its highly wrinkled appearance, and Δ*epsB*, which resulted in a smooth, structureless macrocolony. This suggested that g_epsA, which targets U2701, downregulates both *epsA* and *epsB*, which are co-transcribed. Downregulation of *epsC* (U2699) by targeting the sequences within the EAR or *epsC* resulted in the formation of a completely smooth and viscous macrocolony, similar to that of the Δ*eps* mutant (Figure 2a). This is consistent with the importance of the matrix exopolysaccharide biosynthetic pathway, encoded by *epsC-O*, in the formation of highly wrinkled biofilms, as previously established ^44^.

Downregulation of the bacterial hydrophobin gene *bslA* using g_bslA_2_, which targets *bslA* 69 bp after the start of the ORF, resulted in a smooth macrocolony with a phenotype similar to that of the Δ*bslA* strain (Figure 2b). Interestingly, downregulation of the non-coding S1185 element in the 5’ UTR of *bslA* using g_bslA_1_ did not completely abolish wrinkle formation. These observations underscore the potentially variable efficiency of CRISPRi knockdown, which is contingent on the position and sequence content of gRNA relative to the targeted gene, as previously described ^45^.

Subsequently, we investigated the silencing of the *tapA-sipW-tasA* operon, which encodes TasA-amyloid fiber assembly. Transcriptional studies have demonstrated that the region is traversed by two transcripts, one encompassing *tapA-sipW-tasA* and the other only covering *tasA* (Figure 2c) ^31^. The downregulation of the *tapA-sipW-tasA* operon by *g_tapA* resulted in a macrocolony with impaired wrinkle formation, similar to that of the *ΔtapA-sipW-tasA* knockout. In contrast, the single Δ*tapA* mutant strain displayed a mild yet marked phenotype with respect to the macrocolony structure (Figure 2c). This finding is consistent with the role of SipW, which possesses a peptidase activity in maintaining matrix integrity and three-dimensional (3D) structure of the macrocolony ^9^. Silencing of *tasA* by two different guides, one targeting the intergenic region between *tasA* and *sipW* within the 5’ UTR of *tasA* (g_tasA1) and the other targeting the beginning of the *tasA* ORF (g_tasA2), resulted in distinct phenotypes. The guide g_tasA-1 produced a macrocolony phenotype similar to that of Δ*tapA-sipW-tasA*, potentially due to a reverse polar effect on the upstream part of the operon, as previously observed in *B. subtilis* when targeting a gene within an operon ^46^. Contrastingly, the expression of guide g_tasA-2 still resulted in a wrinkled macrocolony phenotype. Both phenotypes further differed from that of the NDmed Δ*tasA* strain, which exhibited a “spreader” phenotype characterized by a thin,extended macrocolony with a poorly wrinkled central core ^7^. This suggests that *tasA* may be subject to specific regulation within the *tapA-sipW-tasA* operon. However, the phenotypic similarities observed among g_tapA, g_tasA1, and Δ*tapA-sipW-tasA* highlight the synergistic interaction between TapA and TasA. This is consistent with the established chaperone-like function of TapA in assisting in TasA assembly into amyloid fibrils within the biofilm matrix ^8,47^.

#### 2.2. Analysis of the dynamics of biofilm formation

The dynamics of macrocolony and pellicle formation were examined through serial imaging conducted at 30-minute intervals over 48 h (Figure 3, Videos S1 and S2). Measurements of macrocolony areas over time were performed for both KO and KD strains of *epsC, bslA, tasA*, and *tapA*, in comparison to the NDmed strain and the NTC (non-targeting control) strain, which expressed dCas9 and a non-targeting guide RNA. The kinetic profiles of the KD strains were comparable to those of their cognate KO strains (Figure 3a, Figure S2a). Macrocolony area was significantly reduced upon silencing of *epsC*, exhibiting a pattern analogous to that observed in the Δ*eps* strain. While the kinetic profiles of *bslA* KO and KD strains were also similarly affected, the most striking profile was observed for the expanded macrocolony area of both Δ*tasA* and g_tasA-1 KD strains (Figure 3a, Figure S2a). These observations are in agreement with the observed macrocolony morphological phenotypes of KO mutants in other studies and validate the use of CRISPRi knockdowns for the functional phenotyping of biofilm genes ^30,48^.

**Figure 3.**
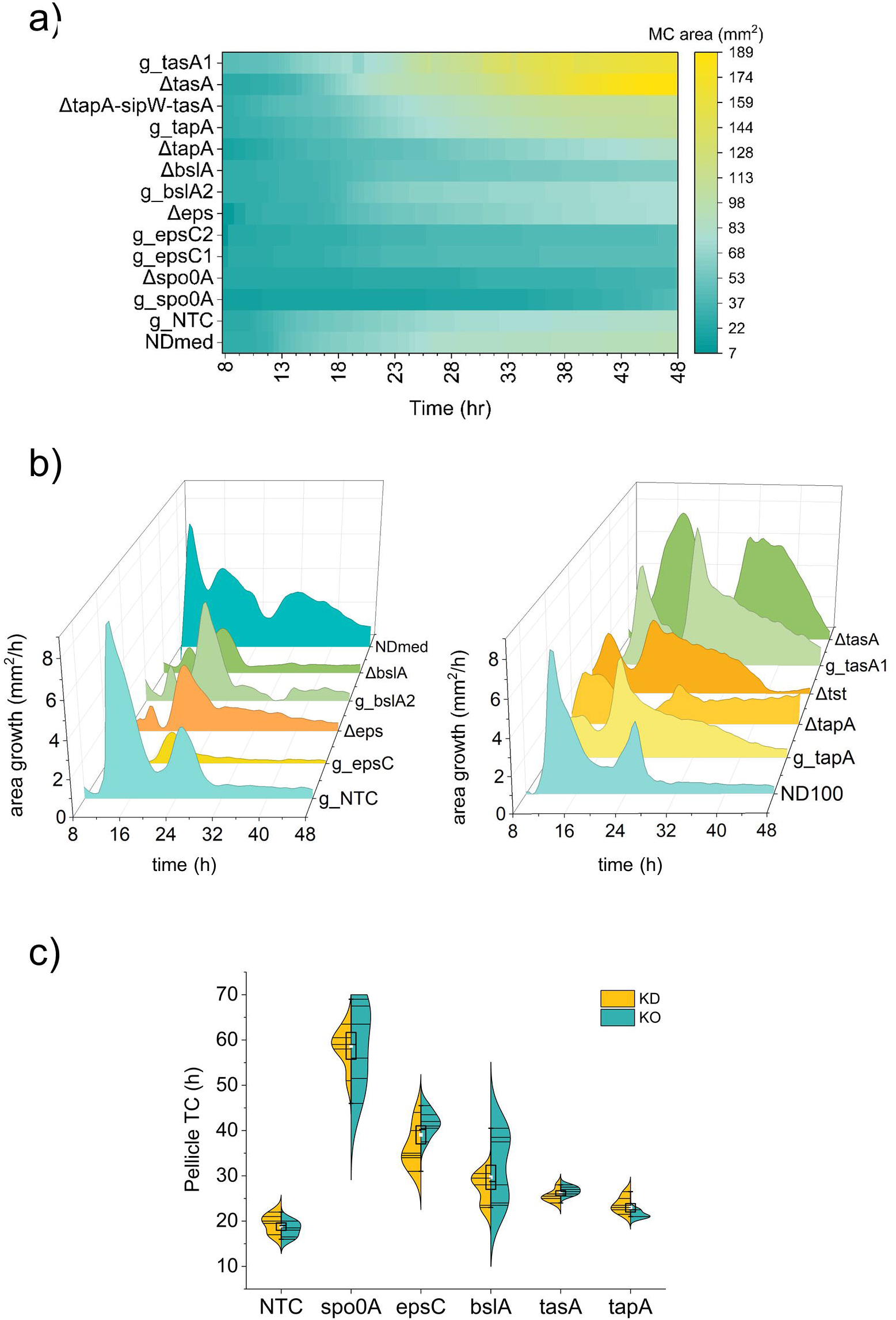
CRISPRi phenotyping of biofilm genes in *B. subtilis* NDmed. CRISPRi-mediated gene knockdowns (g_gene) and corresponding gene knockouts (Δgene) were tested for their ability to form macrocolony biofilms at the solid-air interface (a-b) and the pellicle at the liquid-air interface (c). Δspo0A and g_spo0A strains, which were unable to form biofilms, were used as negative controls. Macrocolony area was measured every 30 min over 48 hours using a BioTek Reshape device. Data represent the mean area and kinetics of area growth (n = 3), displayed in a heat map (a) and a 3D waterfall graph (b), respectively. (c) A split violin plot shows the time at which the pellicle has covered the liquid surface (time of coverage, TC) for wells with gene knockdowns (KD, yellow) and knockouts (KO, blue). Data was derived from n=6 biologically independent static cultures per each strain, with the exception of Δ*tapA* (n

The horizontal growth rate µ of macrocolony expansion was examined to further compare the kinetic profiles (Figure 3b). Throughout the 48 h observation period, the WT NDmed and NDM100 control strains displayed several growth rate phases, characterized by alternating acceleration and deceleration in macrocolony development, forming a wave-like pattern. Notably, the WT NDmed showed three waves, unlike the two waves seen in the NDM100 or g_NTC control strains, which contained the d*cas9* module. This highlights the importance of performing this assay in an isogenic background when comparing KO and KD strains. The depletion of *eps* and *bslA* mostly affected the formation of the first wave, which was highly diminished, while the second wave was present, indicating a strong delay in the formation of the macrocolony (Figure 3b, left panel). CRISPRi-KD of *epsC* exhibited the most severe effect, with a marked decline in the growth rate profile throughout the entire kinetic profile. Silencing of *tasA* resulted in an enhanced growth rate with two larger waves extending up to 48 h, compared to Δ*tasA* and Δ*tapA-tasA* operon, which displayed a largely expanding second wave (Figure 3b, right panel). Analysis of the growth rate of the macrocolony area thus provides valuable insight into potential defects in the formation dynamics. While KO and KD strains are not genetically comparable owing to the inherent operonic effect of the CRISPRi-silencing approach, the macrocolony growth dynamics of *eps* and *bslA* showed similar defects (Figure 3b, left panel). However, the distinct kinetic profiles observed for the *tapA* and *tasA* KO and KD strains underscored the complexity of tapA-sipW-tasA operon regulation (Figure 3b, right panel).

In addition, we compared the time of coverage (TC) at the air-liquid interface, corresponding to the precise time at which the entire surface became covered by the biofilm pellicle, across different sets of KD and KO strains. This assay revealed symmetrical patterns between the KOs and KDs (Figure 3c, Figure S2b, Video S2). A complete pellicle formed between 18 and 20 hours after inoculation for NDmed. While no statistical difference was observed for *tapA* compared to the controls, the TCs were slightly increased for *tasA* and *bslA* and significantly higher for *epsC*, which completed a pellicle between 36 and 40 h after inoculation. Together, these results validated the effectiveness of the CRISPRi approach for investigating biofilm loss-of-function phenotypes at various physical interfaces.

### 3. Competitive exclusion of CRISPRi-knockdown strains in the biofilm pellicle

In the *B. subtilis* pellicle, the interaction between matrix-producing and non-producing cells is characterized by the initial exclusion of the latter from the pellicle. It has been established that *B. subtilis* mutants affected in matrix production, such as *eps* and *tasA* gene knockouts, are outcompeted in co-culture with wild-type (WT) cells and that the matrix remains partially restricted to the matrix-producing subpopulation of cells ^49,50^. We investigated whether non-producer cells resulting from CRISPRi knockdowns also exhibited exclusion from the biofilm. To this end, we built two fluorescent derivatives of NDM100, expressing either GFP or mKate. Using flow cytometry, we investigated the relative abundance of fluorescently labelled cells in pellicles formed by a mixed culture of strains expressing guide RNAs targeting various biofilm genes, in combination with the g_NTC control guide (Figure 4a). First, we targeted spo0A, which encodes the master regulator of cell fate decisions. NDM100 cells expressing g_spo0A were strongly affected in their ability to form pellicles (Figure 3c). NDM100 strains expressing either g_spo0A or g_NTC were tagged with mKate or GFP. Cultures were inoculated at a 1:1 ratio of mKate- and GFP-tagged strains and allowed to form biofilm pellicles. The relative number of fluorescent cells in the pellicle was determined after 24 h of incubation at 30°C using flow cytometry. The pellicles were largely depleted of *spo0A*-silenced cells, with a larger effect when the strains were tagged with mKate than with GFP (Figure 4b). This could be due to the slightly better fitness of NDM100 cells expressing g_NTC when labelled with GFP relative to mKate. Next, the mKate-labelled NDM100 expressing g_NTC was mixed with the GFP-labelled NDM100 strains expressing guides targeting either *epsC* (g_epsC2), *tasA* (g_tasA1), or *bslA* (g_bslA-2). Examination of pellicles after 18 h highlighted the drastic exclusion of *epsC* knockdown cells from the pellicle, as well as the significant exclusion of *tasA* and *bslA* KDs (Figure 4c). These results validated the efficacy of the CRISPRi silencing approach for studying gene fitness in the pellicle biofilm model.

**Figure 4.**
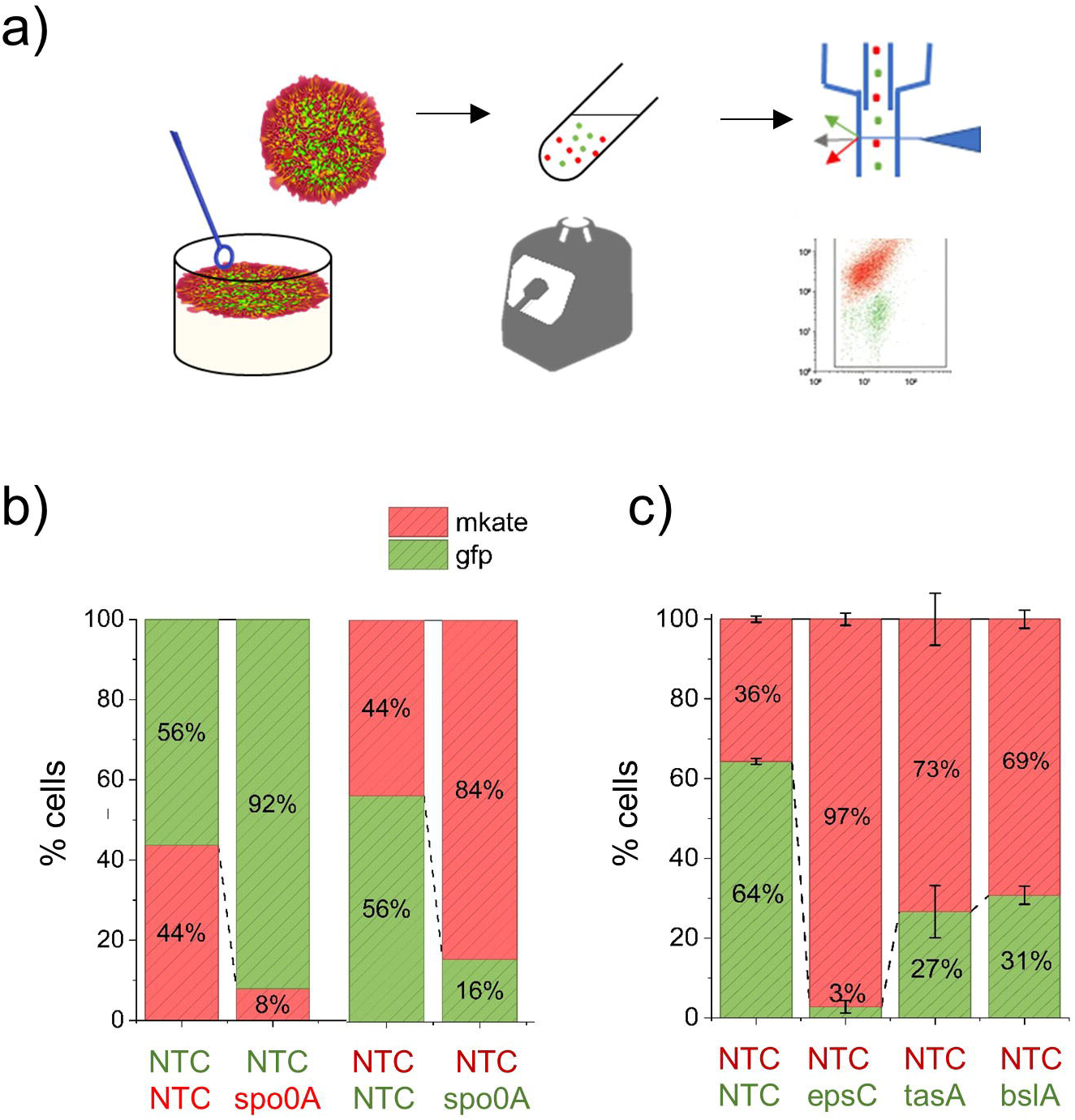
Competition fitness assay in air-liquid pellicles. Experiments were conducted using NDM104 (GFP) or NDM105 (mKate) strains carrying either a non-targeting control (NTC) guide RNA or a guide targeting a biofilm gene. (a) Illustration of the mixed pellicle assay experiment. Green and red fluorescent strains were co-inoculated in the indicated amounts in 12-well culture plates in triplicate and incubated at 30°C under static conditions to form a biofilm pellicle. After 18 or 24 h, the pellicles were harvested, and the cells were extracted by vigorous vortexing. The red-to-green cell ratio was determined by flow cytometry. (b) Fitness of spo0A knockdown in mixed pellicles. The NDM104 strain (GFP-labelled) expressing the NTC guide was mixed with the NDM105 (mKate-labelled) strain expressing a guide targeting *spo0A* and reciprocally at a ratio of 1:1 to form a biofilm pellicle, as described above. The relative proportions of green and red cells in the pellicle were determined using flow cytometry after 24 h. The % values are the averages of the recovered fluorescent cells obtained in triplicate experiments. (c) Competitive fitness of biofilm-gene knockdown strains in a mixed pellicle. NMD105 cells (red) expressing the NTC guide were mixed at a ratio of 1:1 with NDM104 cells (green) expressing a guide targeting biofilm genes (*epsC, bslA*, or *tasA*) to form a biofilm pellicle. The relative proportions of green and red cells in the pellicle were quantified using flow cytometry after 18 h (n=3).

### 4. Involvement of non-coding RNA elements in biofilm formation

Recent transcriptomic data in NDmed identified non-coding RNA (ncRNAs), likely regulatory elements, that were upregulated during biofilm formation ^31,36^. To investigate their potential involvement in biofilms, g_RNAs were designed to silence their expression. These ncRNAs, annotated in *B. subtilis*, were chosen according to their expression profiles during the early steps of biofilm formation (Figure S3a, Table S1). A total of 39 ncRNAs elements exhibiting a fourfold increase in expression (log_2_FC ≥ 2) during the first 7 h were targeted by one or two guide RNAs. After 36 h, the macrocolonies displayed a range of morphotypes, exhibiting distinct patterns based on the presence and configuration of wrinkles (Figure S3b, Figure S4). For the 24 KD strains that exhibited variation in macrocolony phenotypes, the dynamics of macrocolony formation were further examined using automated serial imaging over 60 h (Figure 5, Table S2). The majority of the examined strains (16/24) showed altered dynamics of macrocolony formation upon silencing, with 11 demonstrating a statistically significant increase in growth area over time compared to the NTC control (Figure 5a, Table S2). However, targeting 6 ncRNAs, including BMR26 and BMR28 (BMR is used herein as a shorthand notation for BSU_misc_RNA), which play a role in transcriptional attenuation of the *pyr* operon, as well as BMR10, which is the 5’ mRNA leader of the *pur* operon and corresponds to a purine-sensing riboswitch, associated with S231, a co-transcribed non-coding element, has resulted in a significantly reduced capacity to form macrocolonies (Figure S5a-b, Figure 5a, and Table S2).

**Figure 5.**
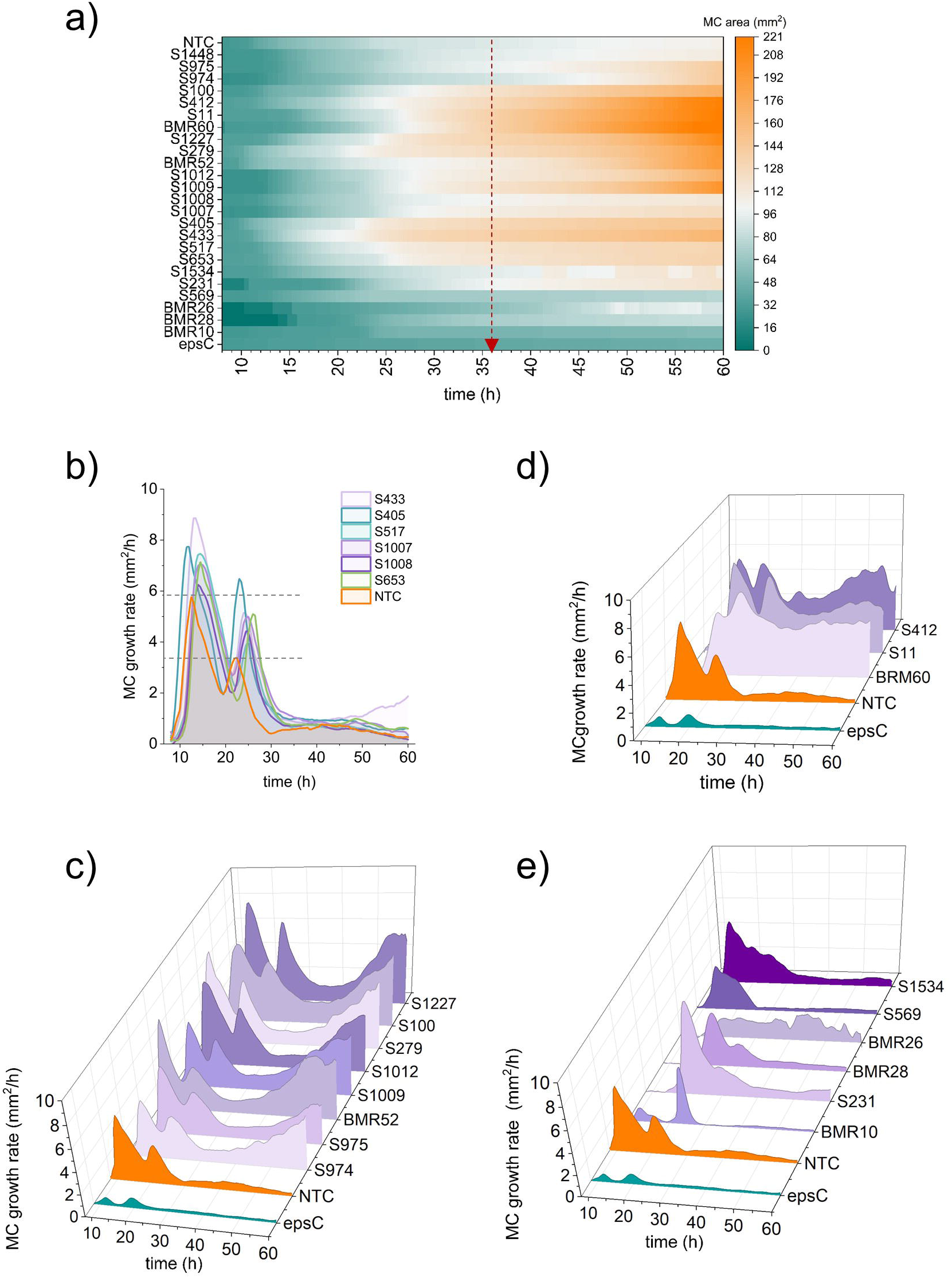
Macrocolony biofilm phenotyping of strains targeting ncRNAs. The kinetics of formation of the macrocolonies were assessed for NDM100 strain derivatives expressing g_RNAs targeting 24 ncRNA elements with marked macrocolony morphotypes. (a) Heatmap showing the kinetics of macrocolony formation over 60 h, as observed using an automated Reshape™ device. The red dashed arrow is drawn at time 36h (b-e) Macrocolony area growth rate (μ) over 60 h. The graphs were organized into four sets of patterns. Data derived from the mean of n=3 independent macrocolonies per strains.

A closer examination of their macrocolony growth rate profiles revealed various behaviors compared to the NTC control (Figure 5b-e). A subset of knockdown strains targeting S653, S517, S405, and S433 exhibited a two-wave growth pattern similar to the NTC strain (Figure 5b). Their expanding phenotype could be explained by a faster growth rate during the first and/or second wave and the persistence of a slightly higher residual growth after 30 h. The second category, comprising seven ncRNA knockdown strains, exhibited two marked first waves of growth, followed by a pronounced additional wave emerging after 40 h. This category was the most represented among the targeted ncRNAs and was exemplified by the *ydaO* riboswitch BMR52, regulated by c-di-AMP, two independent transcripts of unknown function, S1009 and S1227, intergenic transcripts S100 and S1012, and antisense RNAs S974 and S975, potentially counteracting *cwlA*, which encodes a cell wall hydrolase (Figure S5, Figure S6a, Figure S7a). The third category of ncRNAs exhibited the most striking dynamic pattern of macrocolony formation. This category comprises three ncRNAs, namely the riboswitch-like element BMR60, the intergenic RNA regions S11 (located downstream of the cell wall synthesis gene *dacA*, which encodes for a PBP binding protein), and S412 (located upstream of the competence gene *coïA*). In these three cases, the area growth rate did not decrease after the second wave, producing a macrocolony that expanded continuously for up to 60 h (Figure 5d). Finally, in the fourth category, the wave pattern was substantially diminished, explaining the reduced expansion rate of the macrocolonies compared to the NTC control (Figure 5e). In this category, S231 and BMR10 are located in the 5’ untranslated region (5’ UTR) of the *pur* operon, while BMR26 and BMR28 are intergenic elements within the *pyr* operon, highlighting the importance of nucleotide synthesis in macrocolony structure and regulation.

NDM100 formed a highly intricate, wrinkled macrocolony on MSgg medium, similar to NDmed. Downregulation of ncRNAs resulted in various macrocolony morphotypes, displaying distinct levels of wrinkling compared to NDM100 (Figure 6). To capture this morphological complexity, we employed a morphometric approach, which involved the skeletonization of macrocolony images to generate geometric parameters related to the branches and junctions within the complex patterns. A complexity score (C-score) was calculated for each image based on branch number, mean branch length, and junction number of the macrocolony. The C-score was then normalized by the macrocolony area (Figure 6a). With the exception of three strains, all strains showed a significant morphological defect in the macrocolony when compared to NDM100 expressing a g-NTC control guide (Figure 6b). In contrast, targeting S231, BMR28, and S569 resulted in the formation of macrocolonies characterized by pronounced wrinkle formation. These findings indicate that numerous ncRNAs may play a direct role in the development of the intricate wrinkled network within the macrocolony.

**Figure 6.**
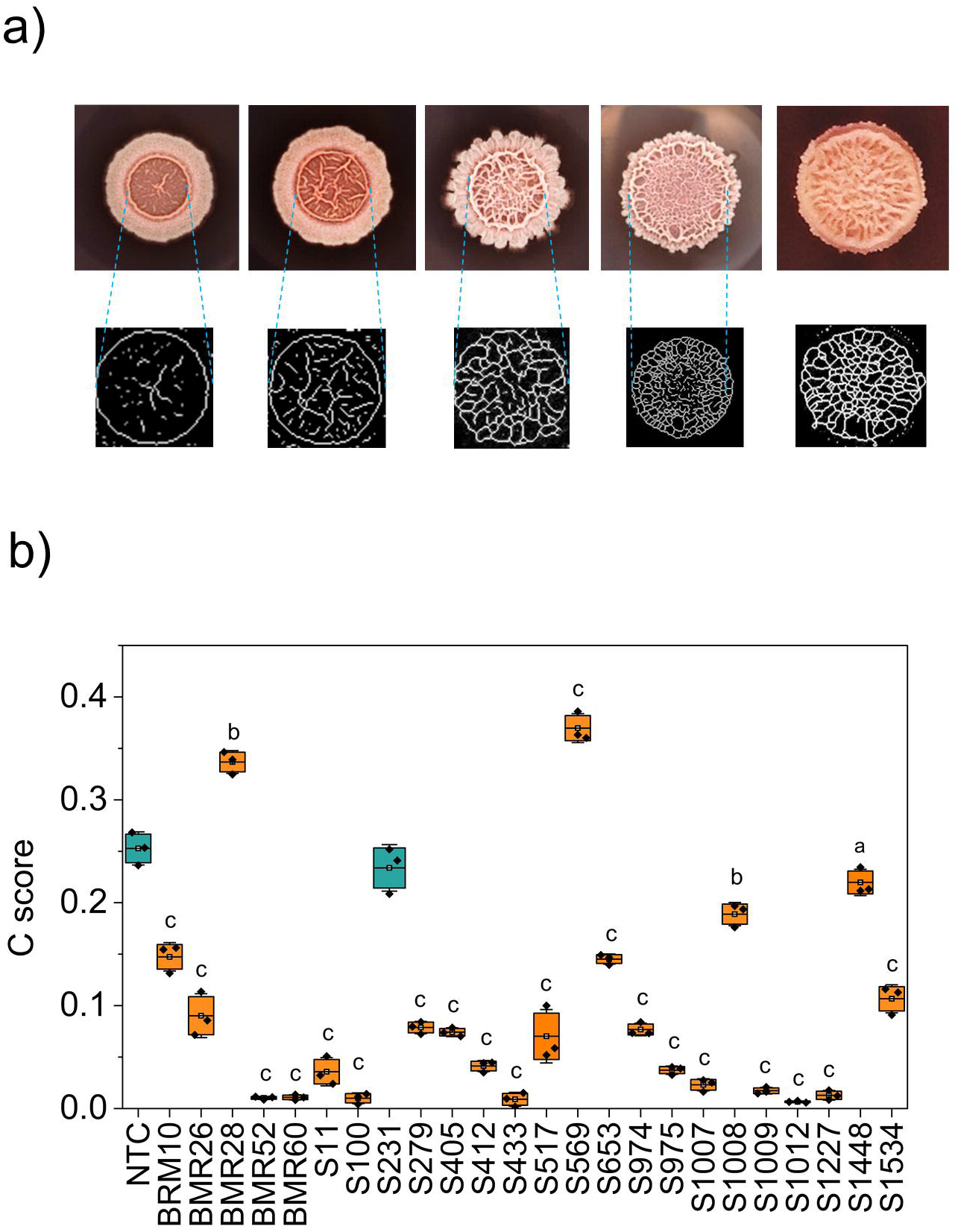
Macrocolony morphotype complexity of ncrNA CRISPRi knockdowns. (a) Determination of a macrocolony complexity score: Images were thresholded to determine the total area of the macrocolony, then adjusted to underline the wrinkles. The number and length of branches and junctions were determined using the Analyze skeleton plugin of Image J ^84,88^. For each macrocolony, a complexity score is calculated, based on the number of branches and junctions and average length of branches, normalized by the total area of the macrocolony (see Materials and Methods). (b) Boxplot of macrocolony complexity scores upon downregulation of ncRNAs. Significant effect is highlighted in orange (a<0.01, b<0.001, c<0.0001; n=3).

We also examined the effects of silencing these ncRNAs on pellicle formation (Figure 7a). A significant delay in TC was observed when 10 of the 24 tested ncRNAs were silenced. Among these were the 5’-UTR elements BMR10, S231 and BMR26 of the *pur* and *pyr* operons. This suggests that targeting ncRNA features involved in nucleotide metabolism pathways reduces the ability to colonize liquid surfaces. Targeting the c-di-AMP-binding riboswitch element BMR52, which controls the expression of the ktrA-ktrB operon involved in potassium homeostasis ^51^, and S517, a ncRNA located downstream of *darB*, which encodes a c-di-AMP-binding protein involved in the stringent response under conditions of potassium starvation ^52^, was also found to delay pellicle formation. This finding underscores the significance of c-di-AMP signaling in pellicle development.

**Figure 7.**
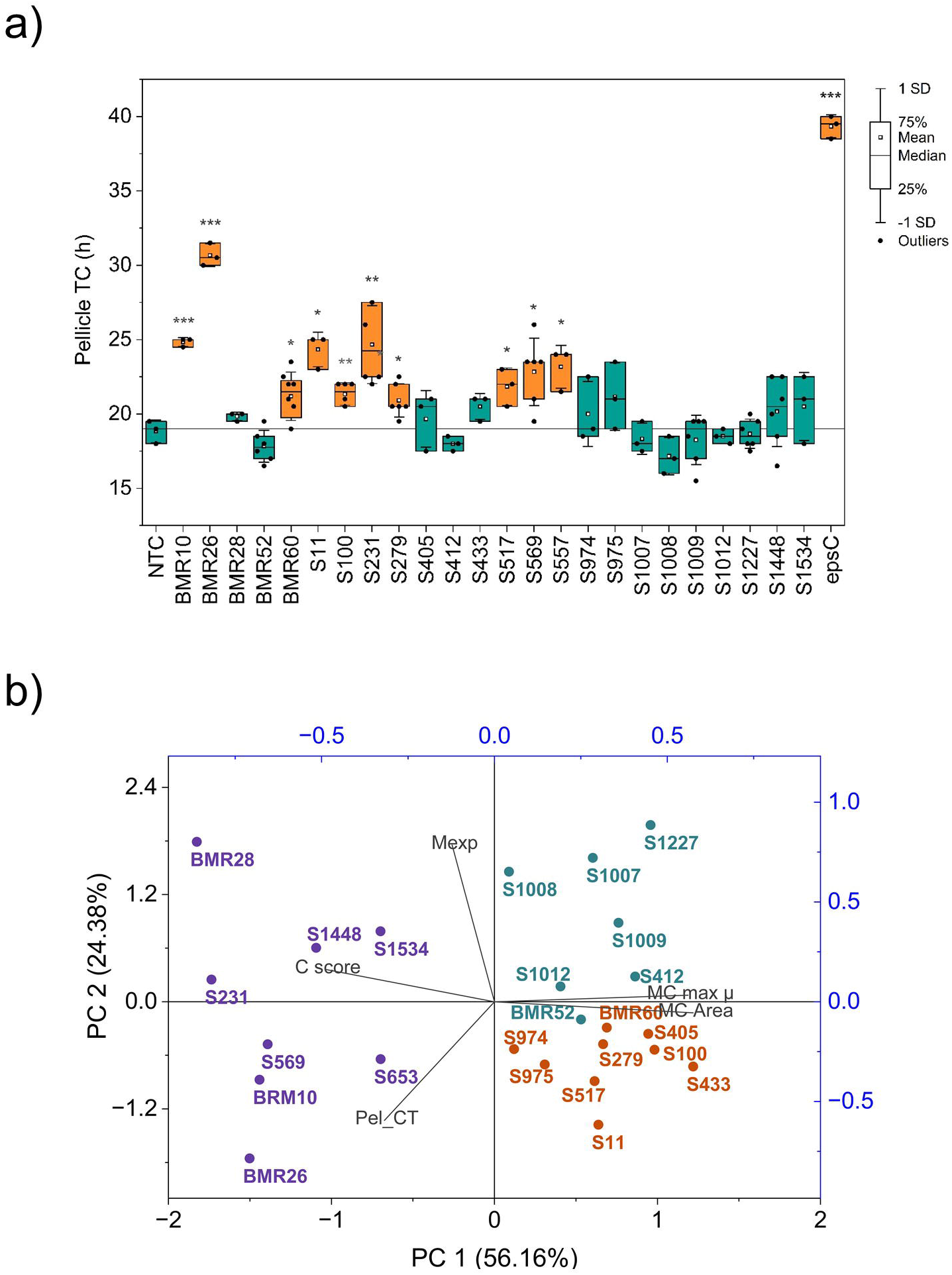
Biofilm pellicle phenotyping of ncRNAs. (a) NDM100 strains expressing a gRNA targeting various ncRNA elements were screened for their ability to form a pellicle at the liquid-air interface. The time of full coverage (TC) of the liquid-air surface by the pellicle was captured using an automated Reshape™ device. The yellow color illustrates a significant increase of TC compared to the control (blue) (* P≤0.05, **P≤0.01, ***P≤0.001, 3≤n≥6). The horizontal line indicates the mean covering time of the NTC control strain. (b) Principal component analysis biplot with hierarchical clustering of biofilm pellicle and macrocolony phenotypes (clusters are indicated by colors: 1 •, 2 •, 3 •). Blue arrows indicate the eigenvectors: Mexp (maximum expression levels during the first seven hours of biofilm formation, as described by Sánchez-Vizuete et al., 2022), Pel TC (pellicle time coverage), MC Area (macrocolony area at 36h), and MC Max µ (macrocolony maximal growth rate) and morphological complexity score (C score). Data relative to this PCA is presented in Table S2.

Principal component analysis (PCA) was conducted on parameters including the pellicle TC, macrocolony area and maximum growth rate, macrocolony morphological complexity score, and expression level of the targeted ncRNA during the first 7 h after the start of biofilm formation (Figure 7b, Table S2). The three principal components accounted for 98% of the variance, with the first two components explaining 56.16% and 24.38% of the variance, respectively. The primary factors contributing to PC1 were macrocolony area phenotypes, whereas the main contributor to PC2 was ncRNAs gene expression (Table S2). Hierarchical clustering analysis (HCA) identified three distinct clusters of phenotypic variation (Table S2). Cluster 1 was largely separated from Clusters 2 and 3 by the first principal component, whereas the second component separated Cluster 2 from Cluster 3. Cluster 1 comprised most of the ncRNAs with strongly altered µ waves (Figure 6e), including BMR26, BMR28, BMR10 and S231 (Figure S5a-b). This observation underscores the role of nucleotide biosynthesis in biofilm formation. Cluster 2 was associated with ncRNA features that are highly expressed during biofilm formation and result in a low-wrinkling and expanded macrocolony phenotype upon silencing. This cluster included a co-located series comprising S1007, S1008, and S1009, which are co-transcribed with *yrpD* a gene of unknown function (Figure S6a). Their location downstream of the *yrpD* coding sequence suggests that this region could be involved in mRNA stability ^31,36^. S974 and S975, the two antisense transcripts opposite *cwlA*, were categorized within cluster 3 (Figure 7b, Figure S7a). *CwlA* represents the terminal gene encoded by the skin element, a latent prophage integrated into the chromosome of *B. subtilis*, which is excised in the mother cell during sporulation ^53^. Although *cwlA* encodes a protein with confirmed hydrolase activity *in vitro*, previous studies have not detected significant expression of this protein in *B. subtilis* strain 168 ^31,54^. However, analysis of the temporal transcriptome under biofilm conditions (see ^36^) indicated very low expression, which was further repressed alongside S974 and S975 during the early biofilm stage (Figure S7b). This observation suggests that these two co-transcribed ncRNAs likely counteract *cwlA* expression. This prompted us to investigate macrocolony formation in a Δ*cwlA* knockout strain, revealing that *cwlA* is important for macrocolony formation in NDmed (Figure S7c-d). While the precise roles of S974 and S975 in the repression of *cwlA* remain to be fully elucidated, these results emphasize the importance of these genetic elements in macrocolony formation. Together, our results highlight the significance of non-coding RNA elements in the development and composition of biofilms on both solid-air and liquid-air interfaces.

### 5. Exploration of gene fitness in biofilm by CRISPRi-seq

#### 5-1 A CRISPRi library targeting genes up-regulated during biofilm formation

As previously demonstrated, using CRISPRi to target key genes associated with matrix production in a mixed population containing matrix producers led to the exclusion of non-producing cells from the pellicle (Figure 4). This finding suggests that the CRISPRi-seq approach, which uses a pool of gRNAs targeting multiple genes, can be applied to investigate gene fitness in biofilm formation on a larger scale. In this type of screening, the reduction of specific gRNAs within the cell population can reveal genes that contribute to bacterial fitness. To this end, we leveraged preexisting data from a temporal transcriptomic dataset in NDmed, conducted during the initial stages of biofilm formation ^36^, and selected genes, non-coding elements, and transcription units (TUs) that exhibited a Log_2_FC ≥ 2 or greater increase in expression during the first 7 h of biofilm formation (Figure 8, Table S3). A collection of gRNAs was designed to target 507 loci, with an average of 4.34 guides per target (Table S3). In addition, 200 guides with random non-targeting sequences were designed to serve as negative controls. A few genes essential for cell survival were also targeted as technical controls. As a result, the library consisted of a pooled population of NDM100 (NDmed CUTE-dCas9) cells, each expressing one of the 2204 single gRNAs. The distribution of read counts after next-generation sequencing (NGS) of sgRNAs showed a slight left-skewed distribution, suggesting that a limited number of guides were underrepresented in the library (Figure S8a).

**Figure 8.**
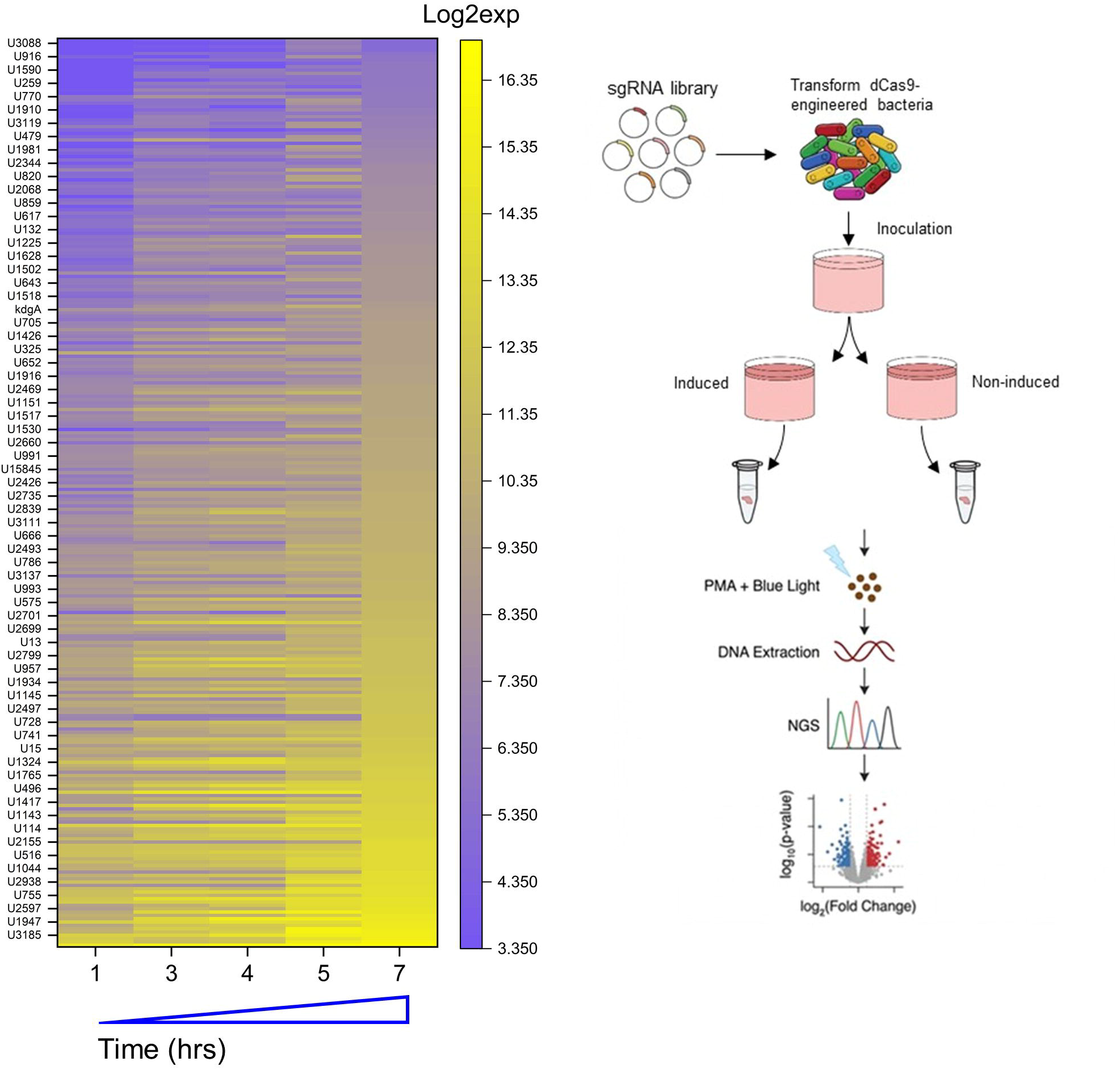
CRISPR pool experimental flow. (Left) Targeted genes were chosen from a temporal scale transcriptomic data performed during early hours (up to 7 h) of biofilm formation^36^. Genes and genetic elements that were upregulated (Log2FC > 2) under biofilm conditions were targeted by an average of five guides. (Right) Experimental workflow for CRISPR-pool in pellicles as described in the material and methods section.

Screens were conducted in rich medium, in the presence and absence of inducers, to identify genes critical for the development of two biofilm models: the biofilm pellicle, at the liquid-air interface, and the macrocolony biofilm, at the agar solid-air interface. Biofilms were harvested after 24 h and dispersed by vigorous vortexing to facilitate cell recovery. Planktonic cells from the liquid phase beneath the biofilm pellicle were also collected and pelleted separately. To prevent PCR amplification from extracellular DNA templates present in the biofilm matrix or from dead cells, all samples were subjected to PMA treatment prior to DNA extraction (Figure 8). The amplicons libraries were subsequently prepared for NGS. Examination of the density plots showing the distribution of all sgRNAs in the respective library revealed a more pronounced negative skewness, indicative of depleted guides, when comparing the induced and non-induced conditions, as well as when comparing the pellicle with the planktonic phase under induced conditions (Figure S8b). For each sgRNA, a fitness score was calculated as a Log2FC comparing its relative abundance between two biological conditions defined by the biofilm fraction and/or induction status of the culture. Gene-level fitness scores were then determined by aggregating the fitness scores of all individual sgRNAs that targeted the same locus. Their significance was evaluated by calculating the q-value from the combination of independent statistical tests for individual sgRNAs targeting the same gene.

The reduction in gene fitness was assessed for each biofilm model. In the pellicle, the differential fitness score for each gene can be evaluated by comparing the relative abundance of guides in the pellicle under induced versus non-induced conditions (PEL_I vs. NI, also defined as PEL) or by comparing the relative abundance of guides in the pellicle with that in the planktonic liquid culture phase under induced conditions (PEL_I vs. PLK_I, also defined as PEL vs. PLK). The fitness of genes in the macrocolony was determined by the relative abundance of gRNA in the presence or absence of inducers (MC_I vs. NI, also defined as MC). Finally, gene fitness was analyzed in the planktonic phase under the pellicle in the presence or absence of cumate and theophylline (PLK_I vs. NI, also defined as PLK). A marked depletion of genes was observed after induction in the different biofilm forms (Figure 9, Tables S4-S8). The analysis of fitness scores identified approximately 54 genes whose gRNA activities resulted in a significant depletion of more than 2-fold (Log2FC < -1, q-valuel < 0.05) from the cell population in at least one of the four models (Figure 9, Figure 10a-b, Table S9). While a specific gene may be depleted in multiple biofilm forms, analysis of its fitness scores can reveal differential depletion levels that may be model-dependent.

**Figure 9.**
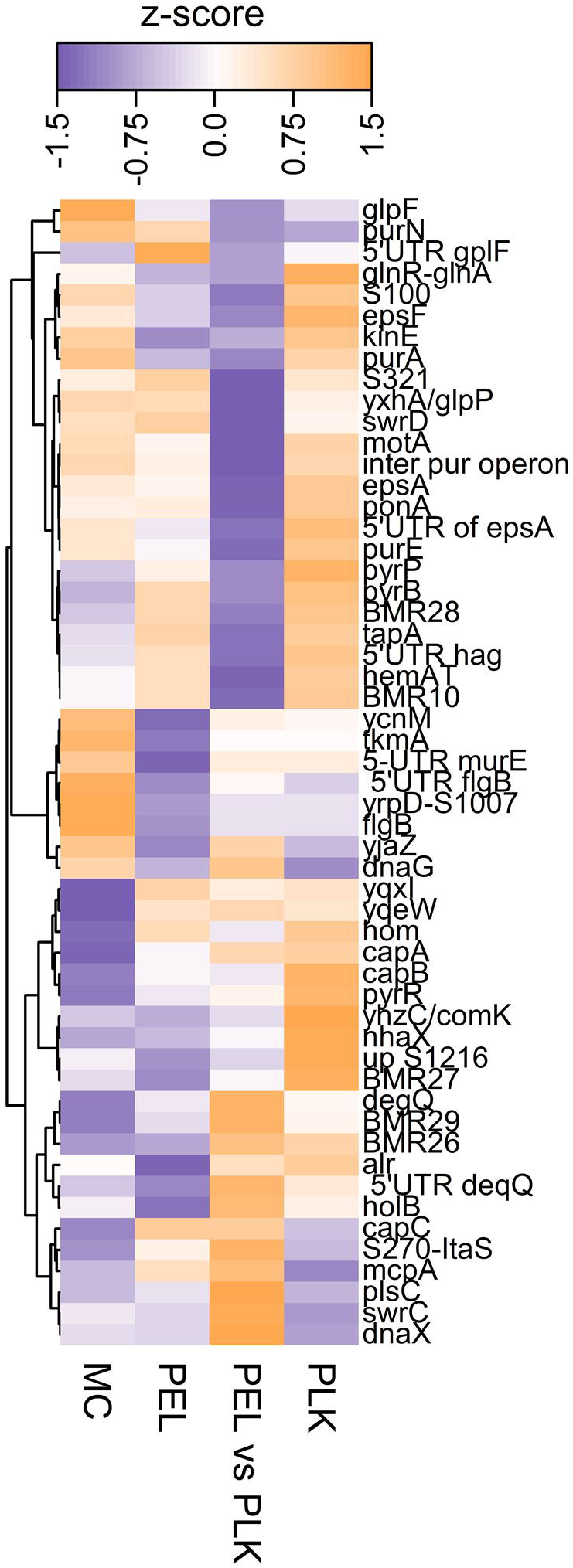
Heatmap with hierarchical cluster analysis of loss-of-fitness genes. The heatmap illustrates the Log2FC values of gene fitness significantly depleted by more than 1.5 fold in at least one model, standardized across the models: Macrocolony (MC, induced vs. non-inducted), Pellicle (PEL, induced vs. non-inducted), Planktonic phase under the pellicle (PLK, induced vs. non-inducted), and Pellicle vs. Planktonic (PEL_PLK, under induced condition). The data and their significance are presented in Table S9. The gene fitness scores were normalized across the models and clustered. Five separate clusters are indicated by the identical colors of the corresponding clusters in the dendrogram.

**Figure 10.**
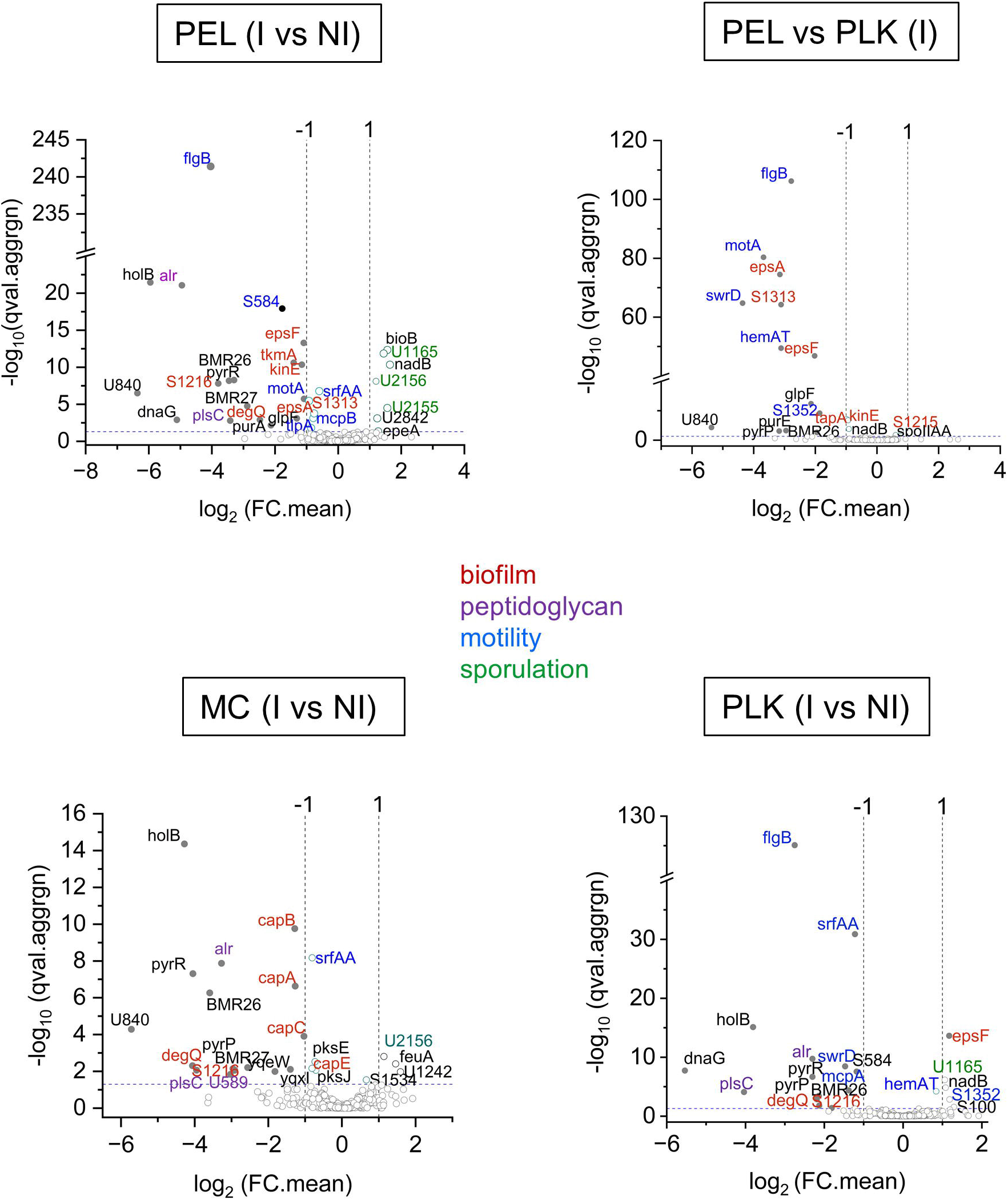
Volcano plots illustrating the loss of fitness genes. Arbitrary cutoff was made for FC >2 to highlight the most significant genes. PEL (I vs NI) model (up left), MC (I vs NI) model (down left), PEL vs. PLK model (up right), PKL (I vs. NI) model (down right). Filled circles indicate genes with statistically significant p-values, otherwise opened. Gene names are colored according to the indicated functional classes.

#### 5-2 Genes required for growth across biofilm models

We first identified a category of KD strains that exhibited a significant reduction in fitness across all models (MC, PEL, and PLK, including PEL vs. PLK). This category likely highlights genes important for growth under these experimental conditions. The first class of genes included *holB, dnaX*, and *dnaG*, which encode integral components of the replisome machinery (Figure 9, Figure 10, and Table S9). The second functional class comprised genes involved in peptidoglycan biosynthesis (*alr*) and phospholipids (*plsC*). These genes are required for cell survival and were initially targeted as positive controls in the pooled CRISPRi library. Notably, targeting *holB* resulted in approximately a threefold increase in severe fitness loss within the PEL, with a reduction exceeding 60-fold in depleted cells under induced conditions compared to the PLK phase (14-fold) and MC (19-fold). Additionally, cells targeting *alr* were depleted by a factor of 30 in PEL, indicating a marked significance of this pathway in this model compared to MC (9.6-fold) and PLK (4.6-fold). Beyond the anticipated decrease in cellular fitness resulting from the suppression of essential genes, this finding underscores the significance of DNA replication and cell wall synthesis in the PEL biofilm model.

Another category of gene targets involved in the purine (*pur*) and pyrimidine (*pyr*) synthesis pathways, was also found to be important. These pathways are not essential for bacterial growth in rich media, although they are important in nutrient-limited medium ^55^. Depleting *pur-*associated genes (*purA, purE, purN*) significantly reduced fitness within the PEL and/or PEL vs. PLK groups. In contrast, knockdown within the *pyr* region ( *pyrR* and *pyrP*) led to a decrease in fitness across all models (Figure 10, Table S9). For most of these genes, a significant fitness loss was also observed in the PEL vs PLK model in the absence of inducers (Figure S8, Table S9). This phenomenon may be attributed to the potential leakiness of the system under the tested conditions. Notably, not all genes identified in our screens exhibited a pre-existing decrease in fitness in the PEL vs. PLK under non-induced conditions. This suggests that these genes may be particularly sensitive to very low levels of basal *dCas9*, which could reflect their importance for the observed phenotypes. Consequently, the presence of a pre-existing fitness defect in the absence of induction is likely to artificially reduce the apparent fitness loss of genes in the PEL and PLK models. Therefore, monitoring significant fitness loss in the PEL vs. PLK model in the absence of induction is likely to underline the importance of the targeted gene in PEL. The biological significance of pyrimidine synthesis pathways in the MC and PEL biofilm models, as well as the PLK model, has already been validated through the analysis of the macrocolony area, morphology, and growth kinetics, along with pellicle formation following CRISPRi targeting of the *pyr*-associated ncRNA attenuator elements BMR26, 27, and 28 (Figures 5-7, Figure S5a). Notably, BMR26 demonstrated an extended pellicle TC of 10 h, compared to the control (Figure 7A). This observation aligns with the 11-fold reduction in BMR26 fitness in the PEL observed in the CRISPRi pool screen, whereas BMR28 was unaffected (Table S9). In our screens, the purine-sensitive riboswitch BMR10, co-transcribed with the downstream ncRNA element S231, *purE*, and *purN*, within the *pur* operon, was found to significantly decrease (up to 7.73-fold) in the pellicle compared to the planktonic phase (PEL vs PLK). *PurA*, which belongs to a different operon, was also found specifically depleted 4.38-fold in the PEL model (Table S9). These data indicate the importance of purine biosynthesis in the PEL model. This was validated by the earlier observation that BMR10 and S231 displayed a ∼ 5h delay in the pellicle CT (Figure 7a). BMR10 and S231 were not identified in the MC model, contrasting with the observation that BMR10, in particular, formed a macrocolony with altered structure and kinetics when targeted (Figure 5e and Figure 6b). Taken together, these findings emphasize the importance of nucleotide synthesis pathways in MC and PEL models.

#### 5-3 Role of peptidoglycan and lipoteichoic acids

In addition to *alr* and *plsC*, other targets implicated in cell wall homeostasis have been identified as contributing to the diminished fitness of KD cells. A moderate yet statistically significant 2.36-fold reduction in cells silencing the U1266 transcription unit, which encompasses the *mur* operon, was observed in the PEL. The *mur* operon is involved in the biosynthesis of peptidoglycan precursors, a pathway which is essential for cell survival. Although this pathway is essential for cell survival, this TU is likely to play a specific role in pellicle formation. The *ponA* gene, which encodes the penicillin-binding protein PBP1 that plays a crucial role in cell division and cell wall morphogenesis, was also found to be depleted by a factor of 2.16 in PEL compared to PLK. Furthermore, the *ltaS* gene, which encodes a major lipoteichoic acid synthase essential for colony development, exhibited a significant fitness loss in both the MC (8-fold) and PEL (4.6-fold). Collectively, these findings underscore the importance of cell wall biosynthesis in solid-air and liquid-air biofilms.

#### 5-4 Differential requirement of eps and cap genes in the pellicle and the macrocolony

The silencing of genes and regulatory elements of the *eps* operon, such as *epsF, epsA*, and S1313, a 5’ UTR RNA element of *epsA*, resulted in the depletion of KD cells, approximately 2-fold, from the PEL population under conditions of *dCas9* induction (Figure 9, Table S9). Depletion reached 4.1- and 8.9-fold, respectively, compared to the PLK phase upon induction, underscoring their importance in pellicle formation. This observation is in agreement with the higher TC of the pellicle previously observed in *eps* KDs and KO (Figure 3C and video 2). Furthermore, this finding was validated by the observed exclusion of *eps* KD cells within the pellicle in the competitive fitness assay (Figure 4b). In contrast, in the PKL phase, cells silencing *epsA, epsF*, and S1313 were enriched 2.2- and 2.9-fold for *epsF* and S1313, respectively. The observed enhanced fitness of *eps* in PLK cells with the already established opposing regulation of motility and biofilm formation ^56,57^.

Unexpectedly, the g_eps guide sequences were not diminished in the MC model (Table S9), which contrasts with the mucoid-type and impaired growth rate patterns observed in the macrocolony of both *eps* KO and KD strains (Figure 2a, video 1, Figure 3a-b). This finding suggests that while exopolysaccharide production is crucial for maintaining the structural integrity and wrinkled morphology of macrocolonies, it may not be indispensable for their formation. In contrast, our findings indicate that γ-polyglutamate production, encoded by the *cap* operon, is a distinctive feature of the MC model. In MC, *capA*, cap*B*, and cap*C* demonstrated a fitness cost of 2.4, which was slightly higher than the 1.8-fold cost observed in PEL and PLK (Table S9). These results emphasize the importance of capsular synthesis in the formation of macrocolony biofilms. To evaluate this hypothesis, we examined the MC phenotype of the *cap* knockout strains. We found that the absence of *cap* genes resulted in smooth macrocolonies devoid of any wrinkled structures (Figure 11a). The three *cap* mutants also exhibited a reduced macrocolony area compared to WT NDmed, and their propagation velocity profiles (µ) were altered in a comparable manner, characterized by a single wave of lower intensity instead of the two distinct waves observed in the WT control strain (Figure 11b-c). These findings underscore the significance of cap genes in the development of solid-air macrocolony biofilms and suggest a potential association between γ-polyglutamate synthesis and the multiphasic growth rate of NDmed macrocolonies.

**Figure 11.**
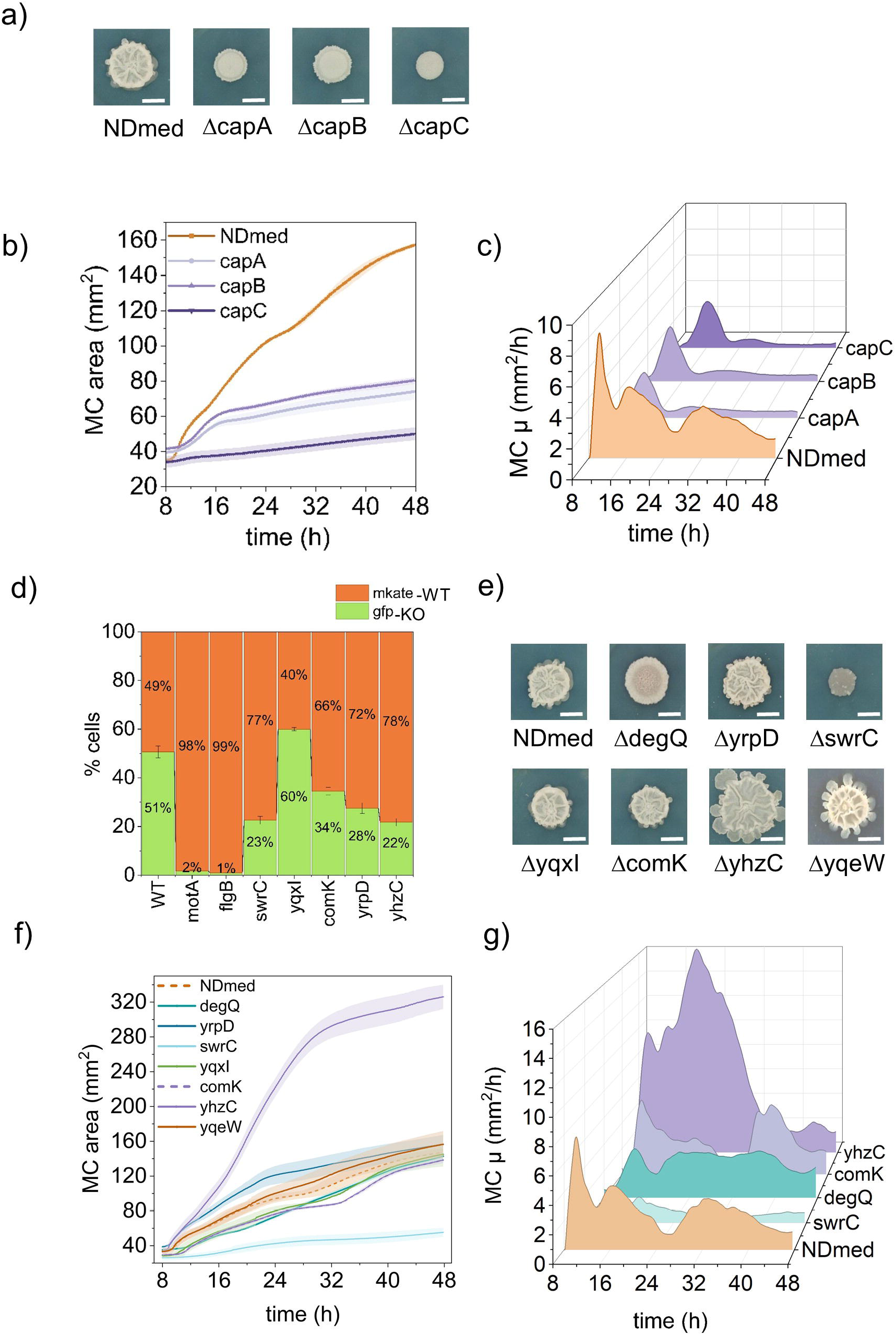
Validation of CRISPRi-screen gene candidates. Cognate gene knockout strains were tested for their macrocolony morphology, area and dynamics of formation (n=3). (a-c) Macrocolony phenotype of gene KOs from the *cap* operon: (a) Macrocolony morphology (a), growth kinetics (b), and growth rates (c). (d-g) Biofilm phenotyping of selected CRISPRi-seq identified genes: (d) Pellicle competition fitness assay. Gene KO constructs were transformed into NDM104 strain to be tagged by the GFP and mixed 1:1 with NDM105 (mKate) prior to inoculation of the static cultures. (e) Morphology phenotyping of macrocolonies (e), area kinetics (f), and growth rates (g) of different gene KOs, as indicated.

#### 5-5 Fitness defect of flagella, and motility gene knockdowns in the pellicle

Depletion of *flgB*, which encodes the flagellar basal-body rod protein essential for flagellar filament assembly, along with its 5’ UTR RNA region S584, resulted in a marked reduction in fitness in pooled library comparisons of PEL and PEL vs. PLK. This was evidenced by 16.4- and 6.8-fold decreases in gRNA abundance, respectively (Table S9). Within the PLK phase, a notable reduction in the fitness of *flgB* was also observed, with a 6.9-fold decrease in sgRNA abundance after induction. This decline in fitness is expected to be reflected in the fitness score of the PEL_I vs. PLK_I model relative to that of the PEL_I vs. NI model. The substantial fitness cost associated with *flgB* highlights the critical role of motility in pellicle formation and the associated planktonic phase. Additionally, the gene encoding the flagellar stator *motA* exhibited a 2-fold decrease in the PEL and a 12-fold decrease in the PEL vs. PLK upon induction (Table S9). The fitness score of *swrD*, which encodes a positive regulator of *motA*, exhibited a 2.8-fold decrease in PLK, while a significant decrease in abundance in PEL vs. PLK was observed under induced (20-fold) and even non-induced (44-fold) conditions (Table S9). These findings underscore the important role of *motA* and *swrD* in pellicle formation. The gene *swrC*, which encodes a surfactin exporter involved in swarming motility, was identified as significant in both the PEL and PLK models, as evidenced by 5- and 7.7-fold reductions upon induction, respectively (Table S9). The importance of flagella and motility genes in pellicle formation was confirmed through a fitness assay, which tracked the relative abundance of KO strains in the pellicle after a 1:1 inoculation of a static culture with WT NDmed and KO strains tagged with mKate and GFP, respectively (Figure 11d). Flow cytometry analysis revealed a major decline in the fitness of *flgB* and *motA*, highlighting their key roles in pellicle formation. Additionally, the importance of *swrC* was highlighted by a decrease in KO cells from 51% to 23% in the mixed pellicle. Surprisingly, Δ*swrC* also exhibited a strong macrocolony defect, whereas *swrC* was not identified as significant in the MC model (Figure 11e). One explanation is that KD cells within the population could benefit from surfactin secreted by other cells in the population. Finally, *srfAA*, another gene in the surfactin synthesis pathway, was also deemed important in all three models, although it exhibited a lower fitness defect ranging from 1.5- to 2-fold. These findings highlight the crucial role of surfactin in the formation of biofilm pellicle, as previously reported for *B. subtilis* ^58^.

#### 5-6 Role of Phosphorylation and chemotaxis

Similar to the *eps* genes, the genes encoding the tyrosine kinase activator *TkmA* and the histidine kinase *KinE* were distinctly associated with pellicle formation (Table S9). This was evidenced by the 2.7 and 2.2-fold decline in sgRNA levels within the pellicle following induction. Additionally, chemotaxis pathways have been identified to play a role in the PEL model. Downregulation of *hemAT*, which encodes a cytoplasmic heme-based transducer responsible for aerotaxis, resulted in a high fitness defect in KD cells, observed when comparing PEL with PLK (8.7-fold), whereas it conferred a fitness advantage in the PLK phase, with a 1.8-fold increase. This observation suggests that cells unable to sense oxygen might be hindered in reaching the air-liquid surface and forming the pellicle, which could explain their enrichment in the PLK phase. Collectively, these results highlight the important role of phosphorylation and aerotactic transduction in pellicle development.

#### 5-7 Silencing of degQ negatively affects its fitness in both the pellicle and the macrocolony

Knockdown of *degQ* was associated with a significant loss of fitness in all models, with a 16-fold depletion of KD cells in the MC and 5- and 4-fold depletion in the PEL and PLK, respectively (Figure 10, Table S9). Targeting S1216, a 5’ UTR ncRNA feature co-transcribed with *degQ*, also led to a fitness decrease in the three models, with the highest reduction taking place in PEL (∼14-fold) compared to MC (∼8-fold) and PLK (3.5-fold). These data highlight the important role of *degQ* in both the pellicle and the macrocolony formation. This was further supported by the examination of the macrocolony area of the NDmed Δ*degQ* strain. Δ*degQ* formed smooth macrocolonies devoid of wrinkled structures (Figure 11e). The kinetics of the expansion of the macrocolony area were slower than those of the WT (Figure 11f). An analysis of the growth rate µ over time revealed a reduced, significantly altered pattern compared to that in the WT NDmed control (Figure 11g). Collectively, these results indicate that *degQ* plays a pivotal role in the formation of these two biofilm forms.

#### 5-8 Importance of glycerol, glutamine and other amino-acid metabolic pathways

In the context of the MC model, *yqeW* and *hom* were identified as particularly significant. These genes exhibited specific fitness defects of 3.5-fold in the MC model (Table S9). *YqeW* is expressed as a monocistronic RNA and encodes a putative transporter involved in phosphate uptake, whereas *hom* is the first ORF co-transcribed with *thrB* and *thrC* from the same operon and is involved in the biosynthesis of the amino acids threonine and methionine. However, our analysis of the Δ*yqeW* mutant strain macrocolony did not reveal significant phenotypic defects compared to WT NDmed, and its role in the formation of the macrocolony remains to be established. The screen also highlighted genes and genetic elements involved in the glycerol uptake and utilization pathway, which were found to be important in all models, although they were particularly important in the PEL model compared to the PLK phase. This was revealed by the 4.4-to-5.7-fold depletion of gRNAs targeting the *glpF* ORF and its 5’ UTR sequences, consistent with other studies that have observed that biofilm formation was stimulated in the presence of glycerol in *B. subtilis* and *P. aeruginosas* ^59,60^. Since glycerol was not added to the rich growth medium used here, the importance of *glpF* in pellicle formation under our experimental conditions was unexpected and might indicate an additional role beyond glycerol transport in biofilms. Finally, a significant fitness loss was observed upon downregulation of the glnR-glnA operon in the three models: MC (5.7-fold), PEL (7.9-fold), and PEL vs. PLK (8.5-fold). This operon is involved in glutamine biosynthesis. In addition to its essential role as an amino acid, glutamine is a major nitrogen source in cells and acts as a signaling molecule for nitrogen availability ^61^. These results suggest the glutamine synthesis pathway plays a key role in biofilm development.

#### 5-9 Unknown genes

Four targets of unknown function were significantly depleted in at least one model. Among these, *yrpD* and *yncM* encode secreted proteins and were found to be specifically depleted in the PEL by 2- and 2.4-fold, respectively. In *B. subtilis*, YrpD and YncM are paralogs belonging to the structural cluster of YrpD-like proteins and are part of the phase transition regulator AbrB regulon ^62^. The *yrpD* gene is co-transcribed with the 3’ UTR series of ncRNA elements S1007, S1008, and S1009. CRISPRi-mediated knockdown of these elements did not affect pellicle formation but did alter the macrocolony structure (Figures 5, 6, 7, and S4). In contrast, biofilm phenotyping of the Δ*yrpD* strain did not reveal morphological alterations in the macrocolony but highlighted a fitness defect in the pellicle when competing with WT cells (Figure 11d-e). This suggests that YrpD may be involved in pellicle formation, whereas the 3’ co-transcribed mRNA regulatory elements, such as S1009, putatively encoding a small protein (25 aa), annotated YrpDX, may play a role in shaping the macrocolony. Analysis of their expression using a temporal transcriptomic dataset under biofilm conditions (see ^36^) revealed that *yrpD* transcription began to increase 4 h after inoculating the static culture (Figure S5c). Although the co-transcribed S1007, S1008, and S1009 RNA elements exhibited a similar temporal dynamic, their detection levels were much lower (∼16-fold), indicating that they are less abundant or possibly more degraded. These observations suggest that *yrpD* and its co-expressed 3’ UTR ncRNAs could functionally diverge, contributing to different aspects of the biofilm architecture.

Among the genes specifically identified in the MC model, *yqxI* attracted our attention because of its genomic localization downstream of the previously identified *clwA*. In the screen, *yqxI* KD cells were depleted by 2.6-fold in the macrocolony. However, the Δ*yqxI* KO strain did not exhibit any defect in the formation or structure of the macrocolony, nor did it display any fitness defects in the pellicle (Figure 11d-e). A possible explanation for this observation is that the guide RNAs (gRNAs) designed to target *yqxI* may exert a reverse polar effect on *clwA*. Finally, our screens identified an uncharacterized small gene, *yhzC*, which encodes a 77 aa-long protein. YhzC was associated with both the MC and PEL models, demonstrating drastic reductions in fitness of 82-fold and 53-fold, respectively, underscoring its critical role in these two models. The genomic location of *yhzC* at the 3’ end of an antisense RNA to *comK* prompted us to investigate the role of these two genes in biofilm formation (Figure S5e). Both Δ*yhzC* and Δ*comK* strains exhibited a marked reduction in fitness, with mutant cells showing a 1.5- and 2.5-fold decrease, respectively, when competing with NDmed WT cells in the pellicle assay (Figure 11d). Furthermore, while the macrocolony phenotype of the Δ*comK* strain remained comparable to that of the WT, the Δ*yhzC* strain developed a macrocolony with a significantly larger area and a distinct wave of growth intensity (Figure 11e-g). These phenotypes confirm the critical role *of yhzC* in the two biofilm models, underscoring its significance as a novel gene in biofilm development in *B. subtilis*.

## 6. Discussion

Bacterial biofilms exhibit significant morphological and environmental diversity. They can develop into various structures, including pellicles at the liquid-air interface, macrocolonies at the solid-air interface, and submerged biofilms. Previous studies have shown that the gene expression profiles of macrocolony and pellicle biofilms are significantly different, reflecting their specialized structural and functional adaptations ^29^. However, our understanding of the specific genetic components or pathways associated with each type of biofilm is limited, which restricts our ability to fully comprehend the molecular basis of biofilm heterogeneity. In this study, we addressed this knowledge gap using a CRISPR interference (CRISPRi) approach to perform extensive functional genomic screening in *B. subtilis* NDmed, a strain known for its genetic tractability and biofilm robustness ^35^. To mitigate potential background effects arising from even minimal levels of leaky *dcas9* expression, we developed a dual-control CRISPR system that ensures stringent repression of *dCas9* at both the transcriptional and translational levels. This system, which integrates cumate-inducible transcriptional control of gene expression with a theophylline riboswitch, was shown to remain tightly closed in the absence of inducers and to be highly dynamic in response to both inducers during exponential growth. We have validated the effectiveness of this system for analyzing diverse biofilm phenotypes. This strategy allowed us to target and repress both coding and non-coding genetic elements, facilitating the identification of critical components essential for biofilm development in *B. subtilis*. Most importantly, it enabled us to create a pooled CRISPRi library of gRNAs stably integrated into the *Bacillus* chromosome. We applied this high-throughput genetic tool to systematically analyze the genetic elements that regulate biofilm formation in two models: the macrocolony biofilm at the solid–air interface (MC) and the pellicle biofilm at the liquid-air interface (PEL).

We used a non-destructive, real-time imaging approach to facilitate a more in-depth characterization of the impact of genes perturbations on the formation dynamics of these distinct biofilms. We first demonstrated the efficiency of CRISPRi-mediated gene knockdown in producing phenotypes comparable to those of corresponding gene knockouts. The analysis of both the structure and dynamics of macrocolonies of strains deficient in major biofilm components, such as exopolysaccharides (EPS) or the hydrophobin (BslA), revealed mirrored phenotypic characteristics in macrocolony morphology and area expansion kinetics. In particular, the discrete kinetic waves exhibited by the non-targeting control strain were similarly affected in both the gene knockdowns and their deleted mutants. Additionally, the time required for total surface coverage by the pellicle (TC) was similarly affected between the knockdown and knockout strains. Our phenotyping analysis of the *tapA-sipW-tasA* operon suggested that the intergenic region between *tapA-sipW* and *tasA* might be under regulatory control. This hypothesis is supported by recent findings in *B. amyloliquefaciens*, which show that this region is subjected to post-transcriptional regulation through a sRNA-sRNA interaction-mediated regulatory cascade that governs the production of amyloid fibers during biofilm development ^63,64^. Overall, our study demonstrated the reliability and relevance of the CRISPRi approach for conducting functional genetic studies on biofilms.

Non-coding RNAs (ncRNAs) are key regulators of various bacterial processes. Although a limited number of ncRNAs have been previously identified to regulate biofilms in *Bacillus*, the large number of ncRNAs encoded within the genome suggests that many more remain to be discovered that potentially play a role in biofilm development. Phenotypic analysis of 40 ncRNAs that were identified as being highly transcriptionally upregulated during the early stage of biofilm development^36^ revealed that their repression often resulted in smooth and/or spreading macrocolonies with decreased wrinkled structures, as illustrated by their low complexity scores (Table S2). Analysis of their expansion dynamics indicated that, for most of them, the increase in area was attributable to altered growth rate profiles. Two distinct patterns emerged: one characterized by a sustained a steady expansion rate after the second wave, which explains the spreading phenotype, and the other is characterized by the reduction or the absence of growth rate waves, resulting in limited macrocolony expansion. (Figure 5). This category comprises ncRNA S11, located downstream of *dacA*, which encodes a PBP enzyme involved in cell wall synthesis, ncRNA S412, located upstream of *coïA*, which is important for genetic competence, and BMR60, a potential guanine-responsive riboswitch element located upstream of *yxkD*, a gene of unknown function. These findings suggest that these pathways are significant contributors to the biofilm formation. The second category includes ncRNAs that regulate the expression of *pur* and *pyr* operons. This finding was further supported by our CRISPRi screens, which identified *pyrR, pyrP*, and *pyrB* from the pyrimidine biosynthesis and acquisition pathway, as well as *purA, purE*, and *purN* from the purine biosynthesis pathway. These results underscore the significance of the nucleotide metabolism in the formation and morphology of *B. subtilis* biofilms. The interplay between nucleotide metabolism and biofilm formation has been reported in other bacteria. Pyrimidine and purine starvation reduce the expression of curli proteins and cellulose in *Escherichia coli*, whereas exopolysaccharide synthesis is inhibited at low intracellular nucleoside concentrations in *Vibrio cholerae* ^65,66^. In *B. subtilis*, the upregulation of purine and pyrimidine biosynthetic pathways has been documented at the transcriptomic, proteomic, and metabolomic levels, during early biofilm growth ^67^. Our findings corroborate the significance of nucleotide availability in the regulation of bacterial biofilm development. However, our data indicate that while pyrimidine metabolism, as evidenced by the genes and non-coding RNAs of the *pyr* operon, was important for both the MC and PEL models, the purine synthesis pathway was significant only in the PEL in our screens (Table S9). Nevertheless, phenotypic analysis of various non-coding RNAs in the *pur* operon revealed their role in macrocolony formation (Figure 5e). The observed difference thus resulted from whether the genetic element is silenced uniformly throughout the entire cell population, as in a KO strain, yielding a macrocolony phenotype, or only in a small subset of cells within the pooled library, leading to population depletion. This issue can be attributed to two factors. First, community complementation in the library may mask the effect of knocking down a specific gene. Second, incomplete silencing of the target gene may leave residual activity, complicating the assessment of its role. Both issues suggest that pooled screening in a structured macrocolony biofilm is likely more challenging than in a pellicle model, as it may obscure the roles of individual genes.

Transitioning to a biofilm state is accompanied by global metabolic remodeling that affects many biosynthetic pathways ^67^. It has recently been shown that nutrient diffusion within the macrocolony generates metabolic co-dependence between the peripheral and interior cells of the colony, leading to oscillations in biofilm growth ^68^. Increased intracellular levels of nucleotides, amino acids, and many other metabolites have been observed in the pellicle during the early stages of biofilm growth, suggesting a coordinated response that supports biofilm-specific processes ^67^. The identification of *glnR-glnA*, involved in glutamine synthesis, and *hom*, involved in the biosynthesis of methionine and threonine, could further illustrates this metabolic reprogramming in the PEL and the MC. However, the significance of glycerol uptake, as illustrated by the identification of *glpF* and its 5’ UTR ncRNA S231, requires further elucidation, as our screenings were carried out in nutrient-rich media lacking glycerol as a carbon source. It is important to note that GlpF is a member of the aquaporin (AQPs) family of proteins in *B. subtilis*. AQPs facilitate passive water flow through the membrane, and *glpF* has been shown to play a role in *B. subtilis* during spore germination ^69^. Although the contribution of *glpF* to PEL and MC remains to be confirmed, our results underscore the potential significance of AQP in biofilm development.

Our pooled CRISPRi screening uncovered various genes and ncRNA elements that displayed differential fitness effects in the biofilm models. Among the genes exhibiting the greatest fold-change reductions, several were identified across all biofilm types. These genes were categorized into functional groups, including DNA replication, nucleotide metabolism, cell wall synthesis, aquaglycerol permeation, and glutamine biosynthesis pathways. Notably, within these general categories, some genes displayed distinctly reduced fitness in a specific model, indicating their potential role in the development of that particular biofilm form. This was observed for *holB* and *alr*, which were 3-fold less represented in the PEL model than in the MC model, whereas the *pyr* pathway was more significant in the MC model (Table S9). Additionally, *yhzC*, a small gene of unknown function, exhibited the lowest fitness score in the screens, with this effect being more pronounced in the PEL model than in the MC. In *B. subtilis, yhzC* is expressed under general stress conditions in a SigB-dependent manner via the production of the polycistronic transcript *yhxD-ascomK-yhzC*, as well as from its own promoter ^70^. Although the role of the anti-sens as-comK RNA in silencing *comK*, which encodes the master regulator of competence ComK, has been demonstrated, the function of *yhzC* remains to be elucidated. The critical role of *yhzC* in both PEL and MC biofilm models was validated by the pellicle fitness assay and the macrocolony formation assay which showed that *yzhC* pays an important role in macrocolony expansion (Figure 11d-g). This outcome highlights the efficacy of the CRISPRi-pool approach in identifying novel key players in biofilm development.

Our screens further identified genes that exhibited effects specific to each of the biofilm models. Regarding extracellular matrix (ECM) components, the importance of *eps* genes, which encode exopolysaccharides (EPS), in biofilm formation has been well-established ^71^. The absence of *eps* genes prevents pellicle formation and significantly alters macrocolony morphology without affecting its formation, as corroborated in this study by the observed effect of the CRISPRi knockdowns. In our CRISPRi-pool screens, we observed differential ECM significance in the two biofilm models, in which the *eps* genes were found to be exclusive to the PEL. In contrast, the biosynthesis of γ-polyglutamate (γ-PGA) was more closely associated with the MC-model. Pellicles require a highly cohesive extracellular matrix to maintain a stable structure at the liquid-air interface, in contrast to macrocolonies, which are mechanically supported by the agar surface. Consequently, the lack of EPS function is expected to cause more severe defects in pellicle formation. γ-PGA primarily contributes to matrix hydration and water retention. These properties are important for both pellicles and macrocolonies but are especially critical for macrocolony development on solid surfaces, where maintaining hydration and three-dimensional structure strongly influence colony morphology. Our results underscore the genetic specificity of macrocolony and pellicle development and morphology, further supporting the relevance of this approach in identifying genes and pathways crucial for biofilm development under varying environmental conditions.

The transmembrane Y-kinase modulator *tkmA* and histidine kinase *kinE* were also found to be specifically involved in PEL. This observation is consistent with the characterized role of TkmA in biofilm formation, presumably through a cross-talk interaction with the non-cognate tyrosine kinase EspB ^24,27^. The role of KinE in PEL formation is supported by its ability to phosphorylate the sporulation and biofilm master regulator Spo0A under biofilm growth conditions ^72^. Most importantly, our findings highlight the crucial role of cellular motility in the PEL model. Genes in the motility pathway, such as *flgB and motA*, were found to be specific to PEL, and their significance was validated in a pellicle competition assay. These findings further support the importance of motility for cells to reach and establish a pellicle at the liquid-air interface ^73^. Additionally, *swrC*, which is involved in surfactin export, was found to be specifically significant in the PEL, in agreement with another study showing that surfactin accelerates pellicle development in *B. subtilis* ^58^. Although found essential for macrocolony formation in our assay, *swrC* was not identified as important in MC in the screen. The capacity of other cells from the CRISPR-pool population to secrete surfactin could account for this observation and suggests that the role of surfactin may differ between the two models. However, the inability of Δ*swrC* cells to form a macrocolony could also be explained by the stress generated by the intracellular accumulation of surfactin, which is suggested to be relieved by extracellular surfactin ^74^. The pleiotropic regulator DegQ was found to be critical in the two models, corroborating its well-documented role in biofilm formation through the stimulation of production of γ-PGA as well as extracellular surfactin and proteases ^75,76^.

Genes of unknown function have been identified in screens as being differentially associated with PEL or MC models. *YrpD* and *yncM*, which encode structurally homologous proteins, were found to be specifically depleted in PEL, whereas *yqeW*, involved in phosphate transport, and *yqxI* demonstrated a loss of fitness in MC. The conserved hypothetical protein YrpD (BSU26820) was annotated as a peptidase_G1_like protein, positively controlled by Spo0A, and repressed by AbrB during vegetative growth. It is thought to participate in controlling the entry into sporulation in *B. subtilis* ^77^. Its selective involvement in pellicle development was validated by the loss of fitness of KO cells observed in the competition assay, wheras no effect of the lack of *yrpD* was observed on the formation or structure of the macrocolony (Figure 11e). In contrast, although the fitness of Δ*yqxI* did not show a fitness defect in the PEL competition assay, its role in macrocolony development requires further validation.

### General considerations from this comparative analysis of CRISPRi screens in biofilms

The identification of multiple genes and non-coding elements within the same genetic unit or pathway provides converging evidence for their critical role in the biofilm model. Notably, significant scores were obtained for genes targeted at the beginning of the ORF, as well as within a co-transcribed 5’ UTR ncRNA region, thereby emphasizing the validity of the gene or pathway. This is exemplified by *degQ* and its associated S1216 and 5’ UTR, *epsA* and its 5 ‘UTR S1313, and *flgB* and its 5’ UTR S584. Furthermore, the results of this study revealed that although g_RNAs are principally designed to target the upstream regions of transcription units, fitness defects can also be observed when targeting genes within operons or intergenic sequences. This suggests that dCas9 can exert its transcriptional blockage effect even far from the loading site of the transcription machinery through potential polar or reverse-polar effects, as previously observed in *B. subtilis* [46, 73]. This is illustrated by the *capABC* and *pyrRPB* operons and their associated intergenic transcriptional attenuators, BMR26 and BMR27.

Finally, it should be noted that while our dually controlled expression system effectively represses the expression of d*cas9* during exponential growth, some limitations have been identified, such as potential leaky *dCas9* expression under biofilm-specific physiological states. Consequently, some targets, presumably highly responsive to even very low levels of *dCas9* from unavoidable leakiness, exhibited a detectable fitness defect when comparing PEL and PLK, even under non-induced conditions. This is expected affect the apparent fitness of the PEL and static PLK models under induced conditions, artificially lowering the measured score. These factors must be considered in data interpretation and experimental design.

## 7 Conclusion

Different biofilm forms exhibit complex spatial transcriptional heterogeneity but differ in their regulatory mechanisms and spatial organization, reflecting their unique biofilm architectures and environmental niches. Importantly, the selective pressures exerted within pellicle and macrocolony biofilms differ substantially, which affects the mutant fitness outcomes in pooled CRISPRi screening. In addition, the identification of non-coding RNAs and regulatory elements highlights the complexity of genetic regulation in biofilm formation, suggesting layers of post-transcriptional control that may fine-tune biofilm development in response to environmental cues. Collectively, these findings demonstrate that biofilm formation in *Bacillus subtilis* is governed by a combination of shared and unique genetic pathways tailored to specific biofilm environments. This dual-model genetic dissection provides a comprehensive view of the molecular basis of biofilm heterogeneity and offers potential targets for modulating biofilm formation in applied settings. This study will also serve as a stepping stone to explore the molecular mechanisms involved in the specific development of biofilms that form at solid-air and liquid-air interfaces.

## 8 Material and methods

### Strains and media

*B. subtilis* strain NDmed is a genetically tractable and highly biofilm-forming *Bacillus* strain originating from a medical device, as described ^35^. Gene knockouts were obtained by transforming NDmed with total genomic DNA extracted from the BKK collection of *B. subtilis* 168, followed by selection with kanamycin. CRISPRi-mediated gene knockdown constructs were obtained by transforming the *B. subtilis* NDM100 strain with pBS1K-gRNA plasmid derivatives, initially created in a commercially available *E. coli* strain NEB® 5-α, for the integration of the gRNA module into the NDM100 chromosome (see below). The plasmids, strains and primers used in this study are listed in Table S10. Strains were cultivated in Luria-Bertani (LB) rich medium supplemented with appropriate antibiotics (ampicillin 100µg/ml or tetracycline 10µg/ml for *E. coli*, and erythromycin 1 µg/ml, chloramphenicol 5 µg/ml, or kanamycin 5 µg/ml for *B. subtilis*. Bacillus biofilms were formed either in Tryptic Soy Broth (TSB) or in in MSgg minimal medium containing 5 mM potassium phosphate (pH 7), 100 mM MOPS (pH 7), 2 mM MgCl_2_, 700 µM CaCl_2_, 50 µM MnCl_2_, 50 µM FeCl_3_, 1 µM ZnCl_2_, 2 µM thiamine, 0.5% glycerol, 0.5% glutamate, 50 µg/mL tryptophan, and 50 µg/mL phenylalanine) ^2,78^.

### Plasmids and constructs

#### Construction of the CUTE-dcas9 expression module for dual expression by cummate and theophylline in B. subtilis (figure S1A)

A theophylline riboswitch aptamer was inserted downstream of the P_veg-CuO_ promoter. To do so, a synthetic DNA fragment carrying the aptamer fused to the 3’ end of the *gfp* coding sequence was exchanged using the restriction enzymes BamHI and PciI within plasmid pCT5-bac2.0 (Addgene # 119872; 10.1007/s00253-018-9485-4). The resulting expression module was composed of *cymR* under the control of the weak constitutive promoter P_xylR_ (P_xylR_-*cymR*) and the gene encoding GFP under the control of the cumate-inducible P_veg-CuO_ promoter coupled with a theophylline “ON” aptamer (P_veg-CuO_ _RBW_theo_-*gfp*) ^38^. This module was subsequently PCR-amplified and transferred between XbaI and PstI sites into the *B. subtilis* Biobrick integrative vector pBS2E ^79^. To enhance *cymR* expression, an exchange between the weak promoter P_xyl_ and the strong promoter P_43_ was performed as follows: To mitigate the potential for growth inhibition that could result from overexpression of *cymR*, as previously observed by Choi et al. ^80^, a low-copy number derivative of pBS2E was first created by swapping the colE1 replication origin to an ori101 origin from the pSC101 plasmid, generating the recipient integrative plasmid pBS2E-ori101. The Pveg_CuO__RBW_theo_-*gfp* fragment was then linked to a P_43_-cymR synthetic fragment by PCR joining, and the resulting module was inserted into the low-copy pBS2E_ori101 vector between the EcoRI and PstI sites, generating pBsCUTE-*gfp*. This system allowed the proper control of expression of *gfp* by cumate and theophylline to be assessed in *E. coli* from the pBsCUTE plasmid as well as in *B. subtilis*, following the insertion of the CUTE-*gfp* module at the *lacA* chromosomal locus. The *dcas9* coding sequence was then fused to P_veg-CuO_ _RBW_theo_ by overlap extension PCR ^81^ to replace the *gfp* gene and generate pBsCUTE-*dcas9*. Furthermore, a translational fusion of the dcas9 gene with *gfp* was created by PCR joining between a PCR-amplified Pveg_CuO__RBW_theo_-*dcas9* and *gfp-*containing fragments, followed by insertion into pBS2E_ori101 to create pBsCUTE-*dcas9-gfp*. The CUTE-dcas9 and CUTE-dcas9-gfp modules were subsequently introduced into the lacA locus of *B. subtilis* NDmed via double homologous recombination, following the transformation of competent cells with linearized plasmids, giving rise to strain NDM100 and NDM101, respectively.

### Construction of the guide RNA module template

A PCR-amplified DNA fragment containing the TrrnB terminator sequence (BBa_K1893035) was inserted between the SpeI and PstI restriction sites of the *B. subtilis* integrative plasmid pBS1C ^79^. The DNA sequence encoding the Cas9 guide-RNA (gRNA) scaffold was placed under the control of the constitutive promoter P*veg* within a synthesized DNA fragment (https://eurofinsgenomics.eu) prior to insertion between the EcoRI and SpeI restriction sites into the pBS1C-trrnB construct This generated the integrative vector pBS1C_P_veg__gRNA-trrnB. To increase the expression levels of *cymR* in *B. subtilis*, the P_xylR_-*cymR-*P_veg-CuO_-*gfp* module from pCT5-bac2.0 was first PCR-amplified using primers PO-01 and PO-04 and inserted into the pBS1C vector within XbaI-PstI. The *gfp* coding sequence was then swapped for gRNA-trrnB by exchanging the sequences between SwaI and PstI, leading to a pBS1C-gRNA module (Pveg_gRNA-trrnB) associated with a copy of the P_xylR_-cymR casette. These constructs were then stably inserted into the *amyE* locus of the *B. subtilis* NDmed genome folloed by delection to chloramphenicol resistance. Later, the gRNA module was inserted into the *amyE*-integrative vector pBS1K ^79^, allowing the selection of *B. subtilis* with kanamycin. Alternatively, the gRNA module was cloned into the replicative plasmid pCT5 between the BamHI and SacI restriction sites.

### Ectopic expression of gRNAs

#### Expression of non-targeting gRNA control-guide sequences from the amyE locus

Two control gRNAs were designed to target a sequence that is not present in the Bacillus subtilis chromosome (non-targeting sequence, NTC). In order to do so, we targeted the synthetic IPTG-inducible HyperSpank promoter and we targeted the 18 base pair (bp) restriction site of the homing endonuclease I-SceI, which is not naturally present in *B. subtilis*. The plasmids pBS1C-gRNA and pBS1K-gRNA were linearized by PCR using primers PO-105 and PO-106, which flank the 20 bp insertion site, and Q5 high-fidelity DNA polymerase. The PCR products were gel-purified and used as double-stranded DNA receptor fragments for the insertion of gRNA targets via the single-stranded bridging DNA assembly method (NEBuilder HiFi). The g_RNA targets were introduced by bridging a 70-bp oligonucleotide containing the 20-bp target sequences, which were flanked by 25-bp sequences homologous to the pBS1K (or C) recipient integration vector (Table S10). The assembly mixture was first transformed into *E. coli* with selection for ampicillin resistance. About 2 to 4 clones were cultivated for plasmid extraction, and the presence of the g_RNA sequence was verified by DNA sequencing. Subsequently, the resulting pBS1C/K-gHpSk and pBS1C/K-gI-SceI were linearized by XhoI prior to the transformation of NDM100 competent cells, followed by selection with chloramphenicol for pBS1C derivatives or kanamycin for pBS1K derivatives. Transformation yielded the insertion of g_RNA modules at the *amyE* ectopic locus of NDM100 by double-crossover homologous recombination.

#### Expression of gRNA targeting guide sequence from the amyE locus

Plasmids pBS1C-gI-SceI or pBS1K-gI-SceI were used as the recipient. The recipient vector was linearized either by PCR using the PO-105 and 106 primer pair as described above, or by digesting by SphI and SapI two restriction sites flanking the 20 bp gRNA target sequence that were inserted in pBS1K-gI-SceI to facilitate the preparation of the linearized vector. The guide sequence was inserted by single-strand bridging DNA assembly using 70 –bp oligonucleotides containing 20-pb targeting a specific gene (Table S10). Plasmids carrying the guide modules were first constructed in *E. coli* prior to being transferred into the NDM100 *amyE* locus as described above.

### Analysis of the dCas9 expression dynamics

#### Analysis of expression at the population level

Overnight cultures of NDM101 cells, carrying the CUTE-dcas9-gfp module inserted at the *lacA* locus, were diluted up to OD_600_= 0.01 and cultivated in LB until reaching an OD_600_ of 0.8-1.0. Cells were inoculated at OD_600_ = 0.1 in 96-well plates containing Luria-Bertani (LB) broth and supplemented with appropriate concentrations of cumate and theophylline. The plates were then incubated overnight at 30°C, under continuous agitation, in a multimodal plate reader (Biotek Synergy 2.0). Both OD600 and green fluorescence were monitored every 10-minutes for 18 h. The strain carrying the untagged CUTE-dCas9 module was used as a control for fluorescent background signals.

#### Analysis of expression at the cell level within the population

Cultures were inoculated as previously described and grown at 30°C in a multiwell plate shaker. Samples were collected every hour to monitor GFP levels in 10,000 cells by flow cytometry (CytoflexS, Beckman Coulter) with excitation at 488 nm.

### Construction of fluorescent strains

Several strains were constructed from NDM100 to express a fluorescent protein (FP), either constitutively or upon IPTG induction. For constitutive expression of the green fluorescent protein (GFP), the *gfp* gene, under the control of the constitutive Pveg promoter, was inserted at the ectopic locus *amyE* of NDM100 following transformation with chromosomal DNA from the *B. subtilis* strain 1A1135 (SG13, https://bgsc.org) and selection for spectinomycin resistance. Similarly, constitutive expression of the red fluorescent mKate protein was achieved by transforming NDM100 with the linearized plasmid SG15, allowing insertion of the Pveg-mkate module at *amyE*. For inducible expression, chromosomal DNA from the *B. subtilis* strains 3A39 and 3A40 (https://bgsc.org), respectively carrying genes encoding the mCherry and the GFP under the control of the IPTG-inducible promoter pHyperSpank, were used to transform the NDM100 strain. All the resulting strains were named as listed in table S10.

NDM100 strain derivatives expressing FPs (constitutively or inducibly) were subsequently transformed with a pCT5 plasmid derivative and maintained in the presence of tetracycline 10 µg/ml. The cells were inoculated in fresh media in the absence or presence of cumate (100 µg/ml) and theophylline (4 mM), either at low (0.05) or high (0.5) OD_600_, to visualize the absence or decrease of fluorescence in the presence of inducers by flow cytometry. The samples were analyzed every hour for 5 h.

### Construction of the Biofilm-centered CRISPRi library in NDmed

The CRISPRi library was specifically designed to assess the fitness contribution of genetic elements potentially involved in biofilm formation. We started by identifying 387 transcribed regions of interest based on available genome-wide transcriptomic data ^31,36^. These regions comprised 270 protein-coding genes, 7 RNA genes, and 110 other transcribed elements. These elements belonged to different types: 5’-UTRs (74), 3’-regions (6), inter- and intra-operonic regions (25), and independent regions (5).

We designed sgRNAs to target these regions of interest using a protocol inspired by van Gestel et al. which employed a combination of custom Perl and R scripts^82^. Specifically, we screened for PAM sequences (NGG) on the template strand and considered the 20 nucleotides upstream of the PAM sequence as the spacer sequence of an sgRNA that would target the non-template strand (i.e., the coding strand for protein-coding genes). We discarded some PAM sequences in cases of multiple targets (occurrence of the same 20-nt sequence upstream of another PAM sequence in the genome) or a high risk of off-target effects, as defined by van Gestel et al. (seed length for off-target analysis: 9-nt; maximum number of off-target hits in the genome: 10)^82^. Furthermore, for poly(G) tracts longer than three nucleotides (nt), corresponding to three or more overlapping PAM sequences, we considered only the first two PAM sequences to avoid overlap between recognition sites of sgRNAs targeting the same genetic element. Starting from the 5⍰ end of the gene, we retained the first five PAM sequences that satisfied these conditions to design the sgRNAs. Some short-transcribed regions contained only four or fewer PAM sequences (27% of the targeted regions, with an average length of 136 bp). This resulted in 1,580 sgRNAs targeting 365 transcribed regions of interest (ROIs). We designed additional sgRNAs for transcribed regions of interest with only four or fewer sgRNAs, or those associated with regions of transcription initiation (transcriptional upshifts in the Nicolas et al., 2012, nomenclature). These were designed to target either strand of the region spanning -50 to +10 nucleotides around the detected transcriptional upshift, provided that they contained PAM sequences that satisfied the aforementioned constraints. This yielded an additional 394 sgRNAs targeting 136 transcriptional upshifts. We added 30 positive and 200 negative controls to our list. Positive controls were designed using the same procedure as for the transcriptional regions of interest, applied to 6 essential genes. Negative controls were designed by randomly shuffling the spacer sequence of 200 of the 1,580 sgRNAs designed to target the transcriptional regions of interest, to generate non-targeting sequences. This resulted in a total of 2,204 (1,580+394+30+200) sgRNAs. Within this library, the sgRNAs targeted a set of 507 genes and transcription units (TUs), either within the transcribed region or the promoter region. An average of 4.34 guides (maximum of five) was designed per transcription unit. The sgRNA sequences and targets are listed in Tables S3 and S10.

A total of 2204 guides were thus synthesized as a pool of 70 bp-long oligonucleotides (https://www.genscript.com/). The insertion into the *E. coli*-replicative and *B. subtilis*-integrative vector pBSK1 was carried out by bridging SphI-SapI linearized pBS1K-g-ISceI double-stranded DNA with a 70 bp single-stranded DNA oligonucleotide pool by NEBuilder HiFi SS-bridging DNA Assembly. The reaction mix was then transformed into high-efficiency chemically competent *E. coli* DH5α cells, and clones were selected on 15 cm-diameter LB plates containing ampicillin. After 15-18h of incubation at 30°C, individual clones containing a single gRNA were harvested from the transformation plates by scraping. Approximately 6x10^5^ colonies were collected, ensuring about 27× coverage of the designed sgRNAs. The suspensions were pooled by centrifugation and subjected to plasmid DNA extraction. After quantification by NanoDrop, the purified plasmid pool was linearized using XhoI prior to transformation into *B. subtilis* NDM100 with kanamycin selection for integration at the *amyE* chromosomal locus. Approximatively 2x10^4^ kanamycin resistant clones were harvested in LB containing 15% glycerol and aliquoted into multiple single-use vials prior to being stored at - 80°C. Viability analysis indicated that each vial contained about

### Analysis of biofilm formation dynamics by real-time imaging

Biofilm formation was analyzed using macrocolony and pellicle formation assays in MSgg medium (1×), adapted from the protocol described by Branda *et al*. ^2^ for *Bacillus subtilis*. Antibiotics were added as required, depending on the strain. All experiments were performed at 30 °C.

#### Pre-culture preparation

Bacterial strains were streaked from glycerol stocks onto agar plates supplemented with the appropriate antibiotic and incubated overnight at 30 °C. A single isolated colony was used to inoculate an overnight culture in LB broth containing the corresponding antibiotic. Overnight cultures were grown aerobically at 30 °C with shaking at 200 rpm. The following morning, the cell density was measured by optical density at 600 nm (OD600), and cultures were diluted into fresh MSgg medium supplemented with FeCl_3_ and CAAFS to an initial OD600 of 0.05. FeCl_3_ was added from a sterile-filtered 50 mM stock solution to a final concentration of 250 µM, and CAAFS was added from a 5% stock solution to a final concentration of 0.025%. Cultures were incubated at 30 °C and monitored until reaching mid-exponential phase (OD600 = 0.2-0.4). When required, induction was performed at this stage by adding theophylline to a final concentration of 4 mM and cumate (4-isopropylbenzoic acid) to a concentration of100 µg/ml. Cultures were then incubated further until they reach OD600 = 1 and subsequently used for macrocolony or pellicle assays.

#### Macrocolony assay

Agar plates were prepared the day before inoculation. A 3% (w/v) agar solution was melted and mixed 1:1 with 2× MSgg medium containing the appropriate supplements, yielding a final agar concentration of 1.5% and 1x MSgg. Supplements were added to obtain final concentrations of 250 µM FeCl_3_, 4 mM theophylline, and 100 µg/mL cumate. For endpoint experiments, Congo Red was added to the medium at a final concentration of 10 µg/mL. Five milliliters of medium were added per well in 6-well plates for endpoint analyses, whereas 2 mL per well were added into 12-well plates for experiments without Congo Red for the Reshape Biotech automated imaging device. The plates were allowed to solidify at room temperature and stored until use. For inoculation, the agar plates were dried for 1.5 h under a laminar flow hood. Macrocolonies were initiated by spotting 5 µL of culture (OD600 = 1) in the center of each well in 6-well plates, or 2 µL in 12-well plates. The plates were left open under the hood until the inoculum droplets were fully absorbed and dried. Macrocolonies were incubated at 30 °C with controlled humidity (70%) in a temperature-controlled incubator for endpoint experiments or incubated within the Reshape Biotech automated device equipped with a camera. Macrocolonies were grown until the desired endpoint, typically 48 or 60 h, and images were captured every 30 min. The macrocolony area was monitored through automated AI-assisted detection, enabling the real-time acquisition of growth parameters.

#### Determination of macrocolony morphological complexity score (C)

Digital images of macrocolonies grown at the surface-agar to air were taken after 36 h. Images were segmented by thresholding in ImageJ to first determine the total area of the macrocolony, and then thresholding was adjusted to underline the wrinkles^83^. The thresholded binary images were subjected to morphological transformation in order to delineate the wrinkles and ridges that composed the macrocolony topology. This was achieved using the Skeletonize (2D/3D) plugin^84^. The number and length of branches and the number of junctions were determined using the AnalyzeSkeleton plugin^84^. The complexity score was determined by intergrating both the topological and morphological parameters. The topology score (T) corresponded to the ratio of the number of junctions (J) to the number of branches (B): 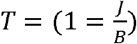 The Morphological score (M) was determined by normalizing the number of branches (B) by their average length (l): *M* = *B* * *l*. The complexity score (C) is calculated by multiplying TxM, normalized by the macrocolony area (A), giving: *C* = [(*B*+ *J*) * *l*]/*A*.

#### Pellicle assay

Pellicle formation was assessed in MSgg medium supplemented with 250 µM FeCl_3_, 4 mM theophylline, and 100 µg/mL cumate. Cultures grown to OD600 = 1 were directly inoculated into the medium at an initial OD600 of 0.05. For endpoint experiments, pellicles were grown in 6-well plates with the addition of Congo Red at 10 µg/mL, whereas experiments in the Reshape Biotech^T^ automated imaging device were performed in 12-well plates without Congo Red. Pellicle cultures were incubated statically at 30 °C in a standard incubator without humidity control or within the Reshape Biotech system and grown for 24 h with image capture every 30 min. The timing of pellicle formation defined as the appearance of a film covering the entire surface of the well (i.e. time of coverage TC), was determined as previously described ^58^.

### CRISPRi-pool screening

#### Library recovery and pre-amplification

An aliquot of the pooled CRISPRi library was retrieved from storage at -80 °C and thawed on ice. The entire volume was diluted 100 fold in freshly prepared Tryptic Soy Broth (TSB) without antibiotics and incubated at 30 °C with shaking for 2 h to allow recovery of the cells. To allow the library to reach exponential growth, cell density was measured by optical density, diluted into fresh TSB to an OD600 of 0.03 and let grow until reaching an OD600 of 0.3. The culture was then allowed to grow for one more doubling time before being split equally between two Erlenmeyer flasks. For the induced condition, one flask was supplemented with theophylline and cumate. The cultures were then incubated at 30 °C with shaking until an OD600 of 0.6 was reached, before to inoculation of the biofilm assays.

#### Macrocolony and pellicle CRISPRi screening

Pellicle and macrocolony screens were performed in triplicate and duplicate, respectively, following the same procedures described for biofilm assays, using 12-well plates for the NGS experiments. For macrocolonies, 2 µL of culture was spotted in the center of each well containing 2 mL of dried TSBA agar prepared as described above, with or without inducers as required. For pellicle formation, cultures were inoculated into 2 mL of liquid TBS medium per well of a 12-well plate at an initial OD600 of 0.05, with or without the addition of inducers. All biofilm samples were incubated statically at 30°C for 24 h.

#### Macrocolony harvesting and PMA treatment

After 24 h, the entire macrocolony was carefully removed from the agar surface using a sterile loop and transferred into a 2 ml Eppendorf tube containing 1 mL of sterile 1xPBS, ensuring complete recovery of the biofilm biomass. Samples were vortexed for 5s followed by a short pause, repeated twice, and then vortexed continuously for 15 min to disperse the cells and disrupt the extracellular matrix. At the end of this step, no visible aggregates were observed. Cells were pelleted by centrifugation at 5,000 × g for 10 mn at room temperature. The pellets were suspended in 400 µL of 1x PB. Propidium monoazide (PMA, Biotium) was added to a final concentration of 50 µM (1 µL of a 20 mM stock solution was added to 400 µL of the cell suspension). The samples were incubated for 10 min at room temperature in the dark on a rotating device, then photo-activated under blue light for 15 min. After photo-activation, the samples were centrifuged at 6,000 × g for 10 min, and the pellets were stored at −20 °C until genomic DNA extraction.

#### Pellicle harvesting and PMA treatment

The pellicles were gently scooped up from the interface using a sterile loop and transferred to 2 ml Eppendorf tubes containing 1 ml of 1xPBS. The samples were vortexed continuously for 15 min to dissociate the biofilm matrix and to release the cells. Cells were pelleted by centrifugation at 5,000 × g for 10 min, suspended in 400 µL of 1xPBS, and treated with PMA as described for macrocolony samples. Following photo-activation and centrifugation, the pellets were stored at −20 °C. To analyze the planktonic fraction below the pellicle, 1 mL of the liquid phase underlying the pellicle was carefully collected using a pipette and cells were collected by centrifugation prior being suspended in 400 µL of 1x PBS, and subjected to PMA treatment as described above. The pellets were stored at −20 °C until genomic DNA extraction.

#### Genomic DNA extraction, NGS sample preparation and sequencing

Genomic DNA was extracted from all samples using the NEB Genomic DNA Purification Kit (NEB, T3010L), following the manufacturer’s instructions. Amplicons were generated using primers PO176_reverse-NGS and PO177_forward-NGS (Table S10), using a low-cycle PCR procedure (<22 cycles) to minimize bias. PCR products were separated by electrophoresis on a 1.5% agarose gel and purified using the NEB gel extraction kit according to the manufacturer’s instructions. Amplicons were generated using NGS quality primers consisting of target-specific sequences, fused to Illumina adapter sequences: 5 ⍰ -ACACTCTTTCCCTACACGACGCTCTTCCGATCTaggcaactgaaacgattcggatcctgt-3⍰, and 5 ⍰-GACTGGAGTTCAGACGTGTGCTCTTCCGATCTgatgatgatggtcgacggcgctatt-3⍰ were used as the forward and reverse primers, respectively. Purified amplicons were quantified using a Qubit fluorometer prior to NGS sequencing by Eurofins Genomics.

#### NGS data processing

The 20-nt spacer sequence tags were extracted from the forward reads at the expected spacer position, corresponding to the 136-156 nt range, after verifying that the flanking sequence matched the corresponded to the expected amplicon sequence (maximum total number of mismatches ≤ 2). The extracted tags were then mapped onto the designed sgRNA library to determine the read counts. Read counts were normalized and compared between pairs of conditions using DESeq2 (v1.48.2)^85^ with default settings, including robust estimation of library size and Wald’s test. Library size estimation was used as a control set, considering the sgRNAs with more than 5 reads in each of the 3 samples. The results obtained for the sgRNAs targeting the same genetic element were aggregated by combining p-values using the weighted Z-method ^86^ and by arithmetic averaging the log2 fold-changes. The false discovery rate was controlled using the Benjamini-Hochberg method ^87^ by computing q-values from the aggregated p-values.

## Supporting information

macrocolonies time series

pellicles time series

## Data availability

The data supporting the findings of this study are available within the article and its Supplementary files.

## Acknowledgements

Financial support was provided by the ESA research projects (OSIP IDEA: I-2021-03383) and by the MICA INRAE Microbiology Department.

## Author contributions

H.H.B.: Investigation, Methodology, Analysis, Data curation, Visualization, Writing -Review and Editing. P.N.: Statistical analysis, Writing -Review and Editing. RB: Conceptualization, Funding acquisition, Supervision, Writing -Review and Editing. MFNG: Conceptualization, Funding acquisition, Supervision, Validation, Writing-Original draft, Review and Editing. All authors approved the final version for submission.

## Competing interests

The authors declare no competing interests.

## Additional information

### Supplement Information

#### Supplementary Figures

Figure S1: Expression and functional validation of dcas9 in B. subtilis NDmed.

Figure S2: CRISPRi phenotyping of biofilm dynamics in *B. subtilis* NDmed

Figure S3: CRISPRi phenotyping of ncRNAs.

Figure S4: CRISPRi phenotyping of macrocolonies from strains depleted for ncRNAs.

Figure S5: Genetic context of selected ncRNAs.

Figure S6: Investigation of the *yrpD* gene and 3’UTR.

Figure S7: Validation of the role *cwlA* and asRNAs in macrocolony biofilm.

Figure S8: Density distribution of raw counts of sgRNA in *B. subtilis* NDmed CRISPR library.

## List Tables

Table S1: ncRNA_guides_targets

Table S2: ncRNA_ biofilm parameters

Table S3: gRNA_location_list

Table S4: PEL I vs NI fitness

Table S5 PEL_I vs PLK_I fitness

Table S6 PLK_I vs NI fitness

Table S7 MC_I vs NI fitness

Table S8 PEL_NI vs PLK_NI fitness

Table S9 mean depleted targets across models

Table S10 genetic material

## Notes

### Competing Interest Statement

The authors have declared no competing interest.

### Summary of Updates

The architecture of Bacillus subtilis biofilms is influenced by the coordinated regulation of cellular specialization, matrix assembly, and metabolism. B. subtilis can form different types of biofilm in diverse physical and chemical environments. Understanding the molecular mechanisms that drive biofilm heterogeneity and adaptation to different environmental niches is crucial for developing more effective strategies to control their formation. In this study, we developed a tightly dual-regulated CRISPR interference (CRISPRi) system and employed multi-scale imaging to investigate the functions of individual genes in two distinct biofilm models: the floating pellicle and the intricate, three-dimensionally structured macrocolony, which develop at the liquid-air and solid-air interfaces, respectively. Our findings validated the CRISPRi approach as a powerful method for studying biofilm development over extended periods and revealed that numerous small non-coding RNAs are involved in regulating biofilm growth dynamics and architecture. The CRISPRi approach was also applied to a pool of 507 genes and transcription units, including protein-coding genes and non-coding RNAs, to screen for cell fitness in these two biofilm models. We discovered that, while both biofilm forms rely on fundamental processes such as cell wall synthesis and nucleotide metabolism, they exhibit different genetic dependencies with regard to matrix composition, motility, and signaling. Exopolysaccharide production, motility, and chemotaxis are crucial for pellicle formation. In contrast, macrocolony development is influenced by γ-polyglutamate synthesis and nutrient acquisition functions. Genes of unknown function were also identified to play a differentially important role in the two biofilm forms. Additionally, the CRISPRi screens revealed further non-coding RNAs regulating biofilm architecture and growth dynamics, adding to the existing layers of post-transcriptional control. Collectively, these results demonstrate that biofilm formation at different physical interfaces is governed by a combination of shared and unique genetic pathways tailored to the specific biofilm environment, thereby opening research avenues into the molecular mechanisms specific to the solid-air and liquid-air interfaces.

