## supplementary material for "CRISPR interference functional genomics of coding and non-coding determinants of *Bacillus subtilis* biofilms"

### Supplementary figures :

Figure S1: Expression and functional validation of dcas9 in *B. subtilis* NDmed

Figure S2: CRISPRi phenotyping of biofilm dynamics in *B. subtilis* NDmed

Figure S3: CRISPRi phenotyping of ncRNAs

Figure S4: CRISPRi phenotyping of macrocolonies from strains depleted for ncRNAs

Figure S5: Genetic context of selected ncRNAs

Figure S6: Investigation of the *yrpD* gene and 3'UTR

Figure S7: Validation of the role *cwlA* and asRNAs in macrocolony biofilm

Figure S8: Density distribution of raw counts of sgRNA in *B. subtilis* NDmed CRISPR library

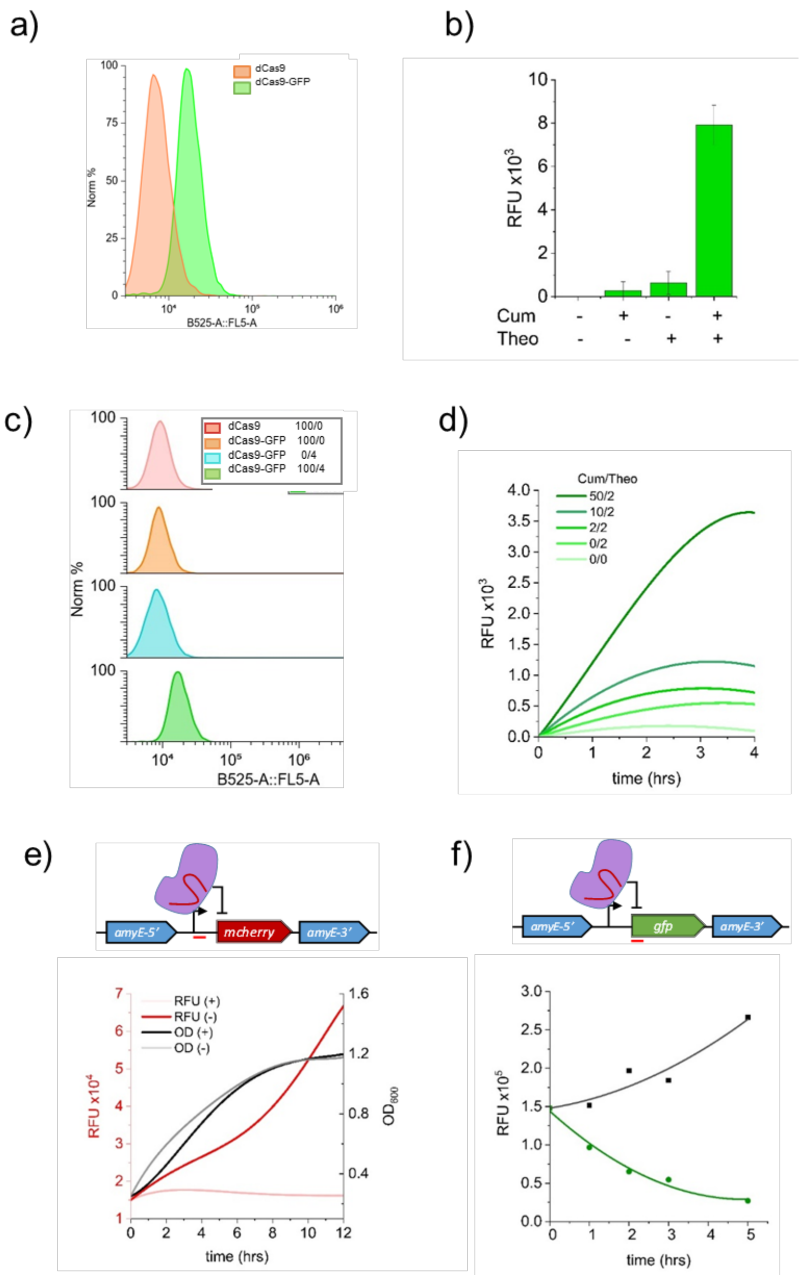


**Figure S1: Expression and functional validation of dcas9 in B. subtilis NDmed.** (**a-b**) Characterization of the CRISPRi system at the cell population level using flow cytometry. NDM100 (*dcas9*) and NDM101 (*dcas9-gfp*) strains were inoculated in the presence of cumate (100µg/ml) and theophylline (4mM). Samples were monitored from $10^4$ cells six hours after the addition of the inducers. NDM100 was used here as a negative control for fluorescence. (**a**) Flow cytometry distribution. Under inducing conditions, fluorescence was distributed within a narrow range across the population, consistent with homogeneous expression. (**b**) Distribution of fluorescence intensities in NDM101 (dCas9-GFP). (**c-d**) Robustness of the dual cumate/theophylline repression/induction system. (**c**) A shift in fluorescence intensity was observed for dCas9-GFP only in the presence of inducers, indicating tight control. (**d**) Early dynamic of expression monitored by flow cytometry every hour for 4 h in the presence of increasing concentrations of cumate (up to 50µg/ml) and/or fixed concentration of theophylline (2 mM). (**e-f**) CRISPRi-mediated repression of fluorescent reporters using the CUTE system. The guide RNAs targeting the *mcherry* hyperspank promoter sequence (g_hpsk) and *gfp* gene coding sequence (g_gfp) were expressed from a replicative plasmid. (**e**) Time course of mCherry fluorescence measured by plate reader following induction *dcas9.* NDM103 strain (mCherry) expressing g_hpsk was inoculated at a low OD (0.05) in the presence (+) or absence (-) of cumate (100 µg/ml) and theophylline (4mM). Culture growth (gray and black) and red fluorescence (light and dark red) were monitored every 10 min over 12 h using a multimodal plate reader. (**f**) Analysis of *gfp* repression measured by flow cytometry. NDM102 strain (GFP) expressing a g_RNA targeting the *gfp* gene was grown to mid-exponential phase OD (0.5) prior to the addition of cumate (100 µg/ml) and theophylline (4mM). Induced condition (green). Non-inducted control (black ). The average green fluorescence of $10^4$ cells was monitored every hour for up to 5 h using flow cytometry.

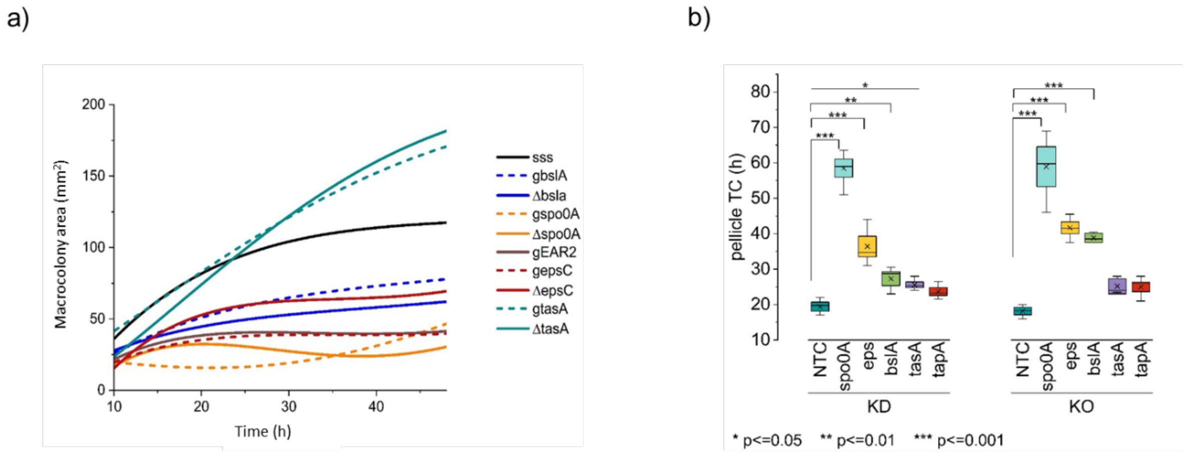


**Figure S2: CRISPRi phenotyping of biofilm dynamics in *B. subtilis* NDmed.** Supplement to Figure 3. (**a**) Macrocolony area kinetics of KD (dashed) and KO (plain) strains targeting genes involved in biofilm formation in *B. subtilis*. Data represent the mean area and kinetics of area growth (n = 3). (**b**) Comparative analysis of the pellicle coverage time of KDs and KOs strains as indicated. Data was derived from n=6 biologically independent static cultures per each strain, with the exception of ∆tapA (n=3).

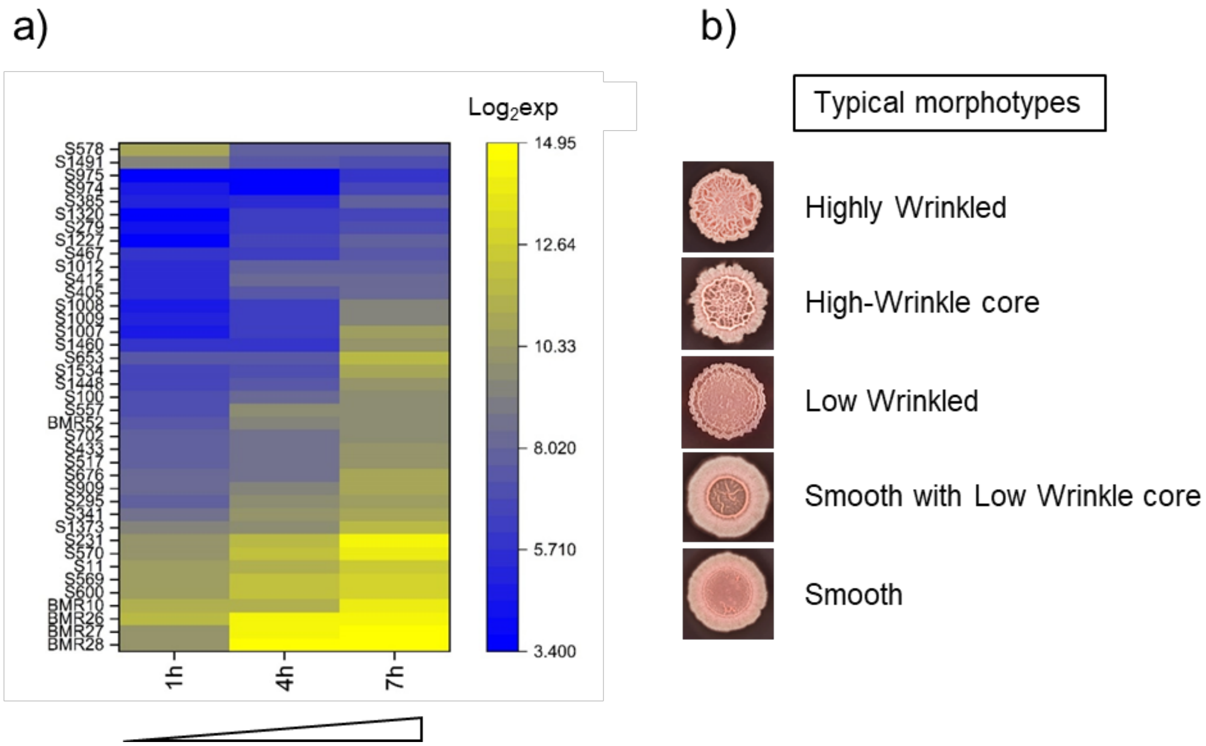


**Figure S3: CRISPRi phenotyping of ncRNAs.** (**a**) ncRNAs were chosen from a temporal scale transcriptomic data performed during early hours (up to 7 h) of biofilm formation. The heatmap illustrates the expression levels of 40 ncRNAs. (**b**)Typical macrocolony morphotypes obtained through phenotyping of the 40 ncRNAs.

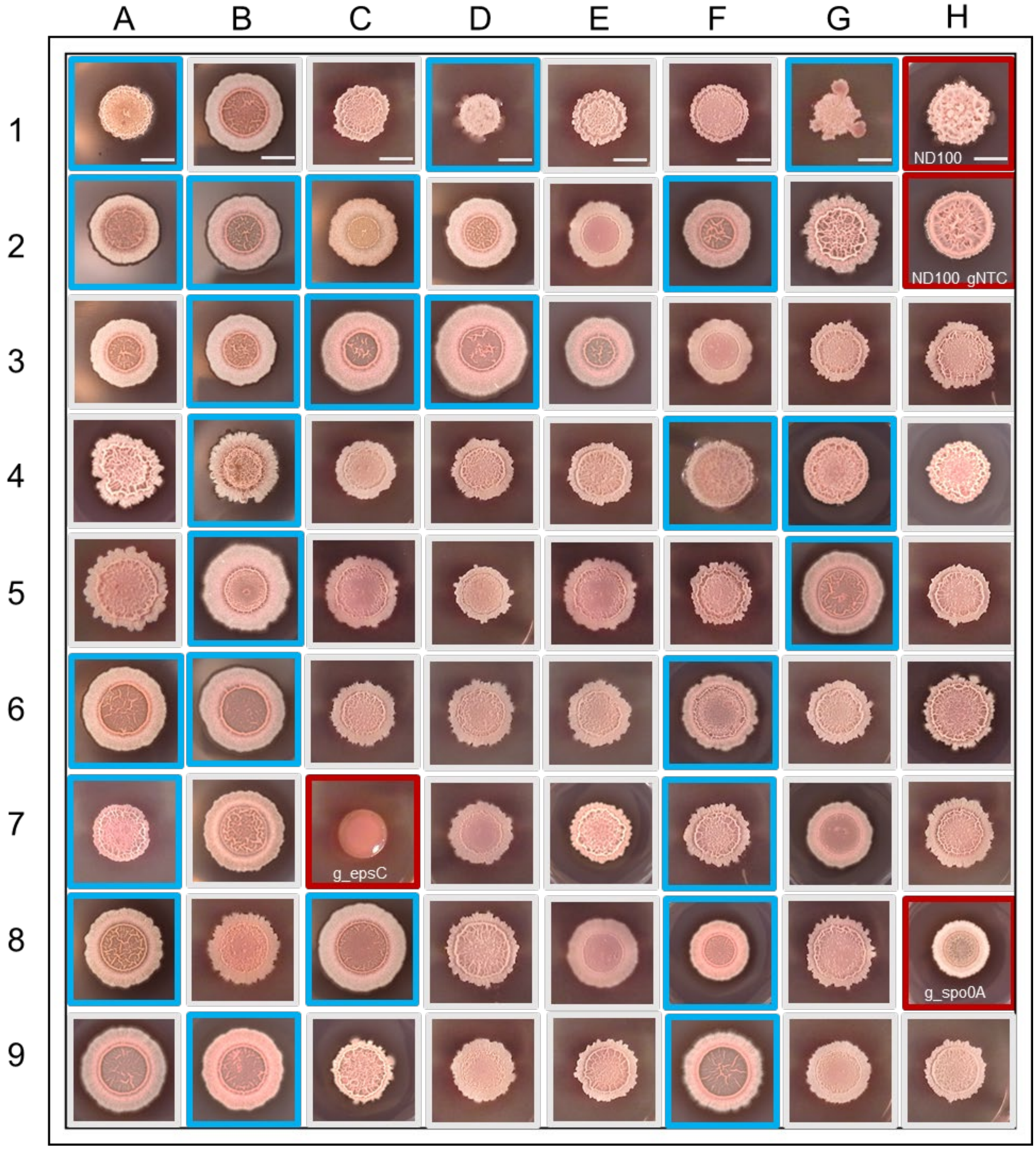


**Figure S4: CRISPRi phenotyping of macrocolonies from strains depleted for ncRNAs.** Gallery of macrocolony morphologies obtained after 36 h in the presence of congo-red. Corresponding ncRNA KDs are listed in Table S1.

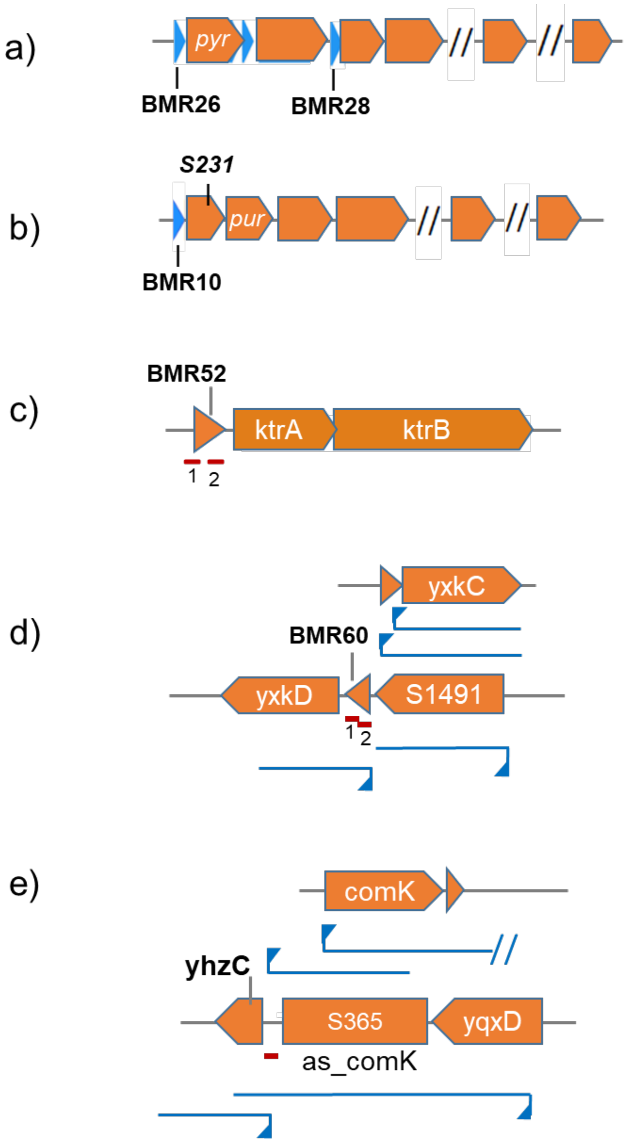


**Figure S5: Genetic context of selected ncRNAs**. gRNA positions are indicated in red. Transcription units, as mapped in Nicolas et al. (2012), are indicated in blue. (**a**) *pyr* operon, (**b**) *pur* operon, (**c**) *ktr*AB operon, (**d**) BRM60 genomic context, (**e**) *yhzC_comK* genomic context.

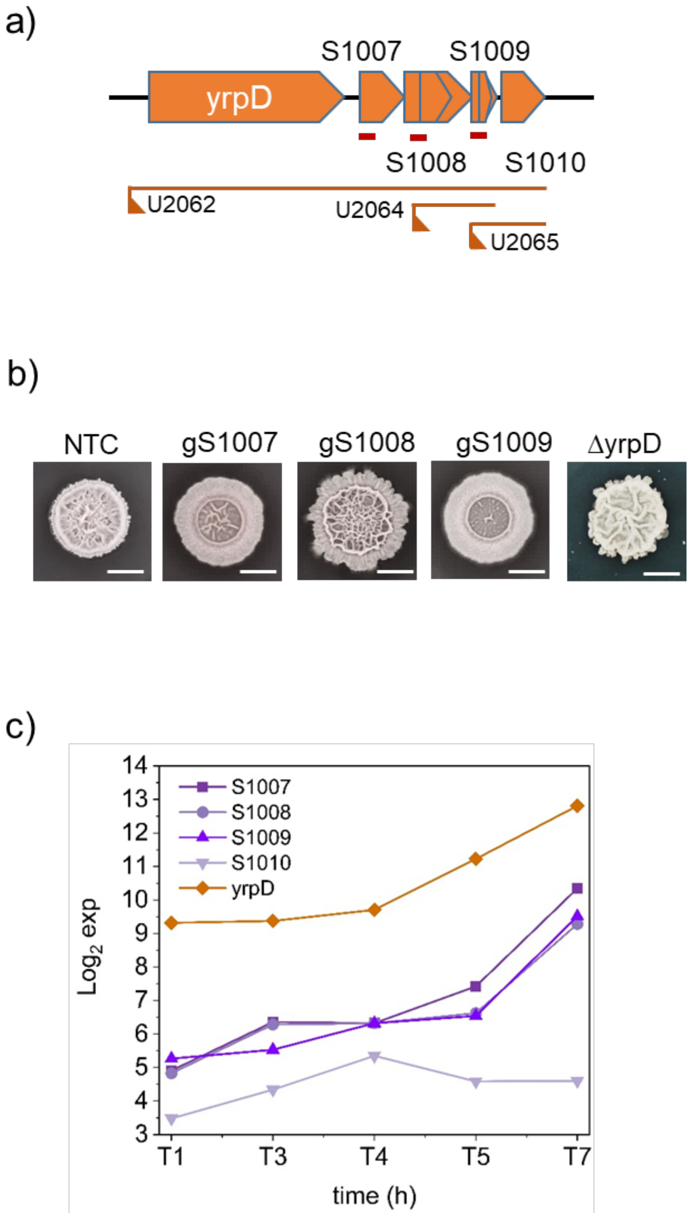


**Figure S6: Investigation of the *yrpD* gene and 3'UTR.** (**a**) Depiction of the genomic context. gRNA positions are indicated in red. Transcription units, as mapped in Nicolas et al. (2012), are indicated in orange. (**b**) Macrocolony morphologies of *yrpD* KO and S1007, S1008 and S1009 KDs. (**c**) Time course expression of yrpD gene and S1007, S1008, S1009 and S1010 3'UTR ncRNA elements showing parallel but not comparable levels of expression.

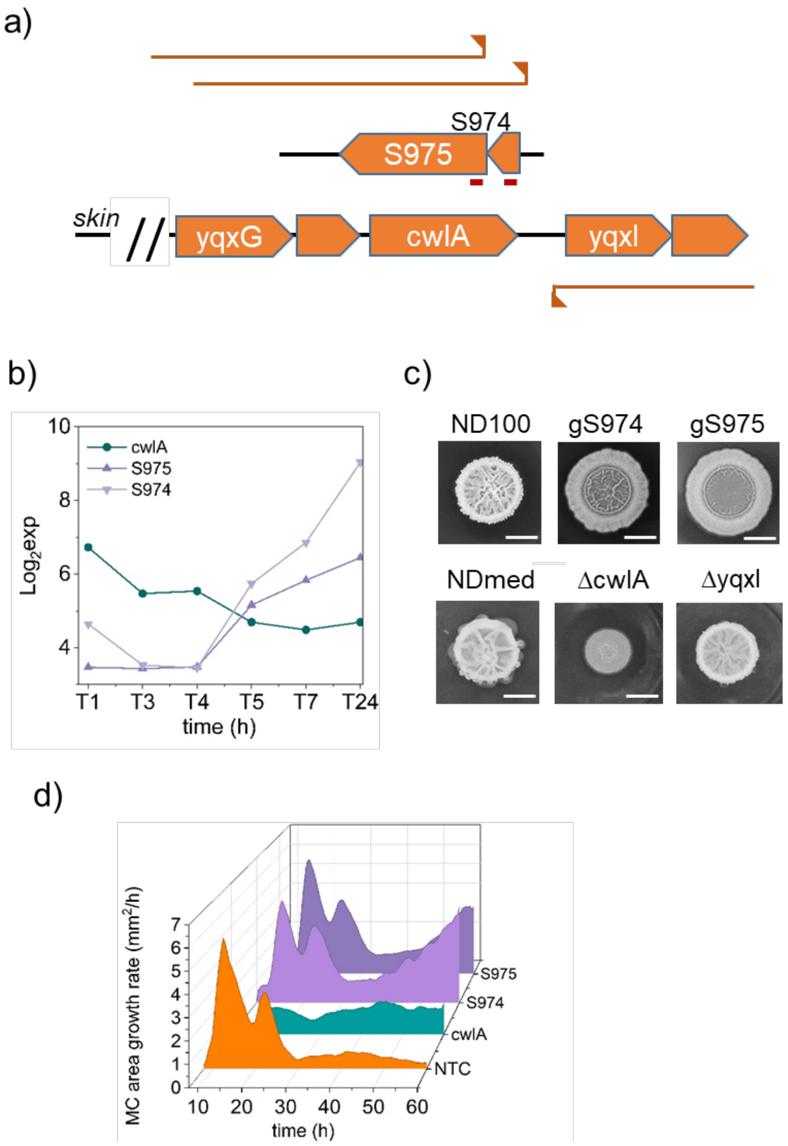


**Figure S7: Validation of the role *cwlA* and asRNAs in macrocolony biofilm**. (**a**) Depiction of the genomic context. gRNA positions are indicated in red. Transcription units, as mapped in Nicolas et al. (2012), are indicated in orange. (**b**) Temporal expression as monitored by Sanchez-Vizuet et al. (2022), showing the inverse correlation of expression of *cwlA* and counter transcripts S974 and S975. (**c**) Macrocolony morphologies of *cwlA* and *yqxI* KOs and S974/S975 KDs. (**d**) effect of *cwlA* and S974 and S975 depletion on macrocolony growth rates. Data represent the mean kinetics of area growth (n = 3).

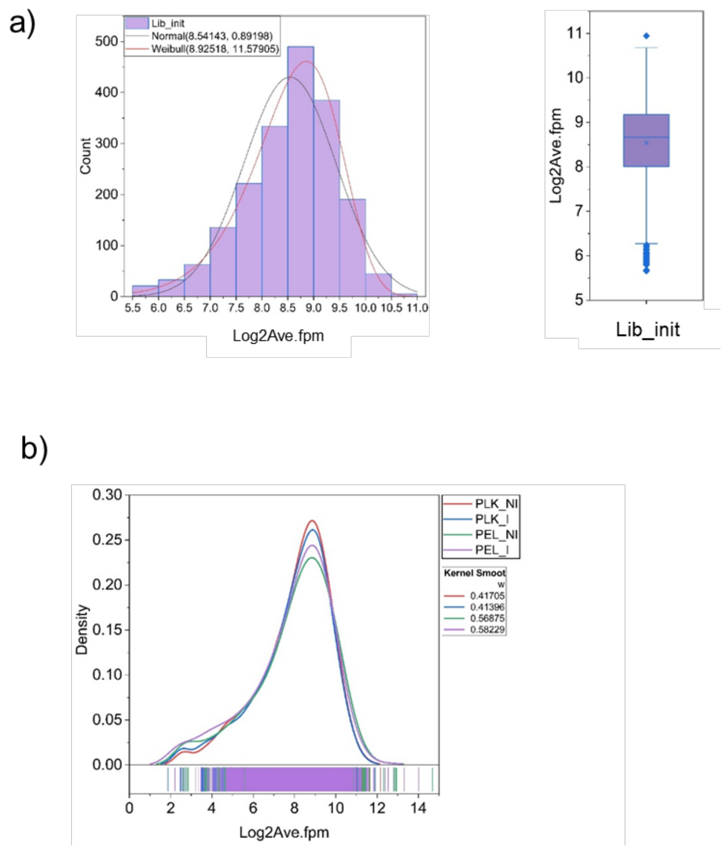


**Figure S8: Density distribution of raw counts of sgRNA in *B. subtilis* NDmed CRISPR library.** (**a**) Density-rug plots showing the distribution of all sgRNAs. Each rug represents one sgRNA. The distribution fits a Weibull shape slightly left-tailed, indicative of the presence of a few outliers with lower abundance in the initial library (skewness 0,64). (**b**) Density of distribution of sgRNA in the pellicle and planktonic liquid phase in the presence or absence of induction. The density-rug plots show the distribution of all sgRNAs. Each rug represents one sgRNA. The left tail shift is indicative of the depletion of guides consequent to the induction of *dcas*9.

### Supplementary tables :

**Table S1: ncRNA_guides_targets. A**- List and DNA sequences of arrayed ncRNA targeted for macrocolony assay. The ncRNA targeted strains selected for further phenotypic analysis using the Reshape are highlighted in blue. **B**- ncRNA chromosomic location, category and expression levels (extracted from Sanchez-Vizuete et al., 2022).

**Table S2: ncRNA_ biofilm parameters**. **A**- Biofilm parameters. Data derived from the mean of independent macrocolonies with corresponding P values and fold change compared to the NDM100-g_NTC control strain: (i) the maximum expression ratio during the first 7 hours (see Table 1B)); (ii) the time of full coverage (TC) of the pellicle at the liquid-air interface as shown in figure 7a (3≤n≥6); (iii) the area of the macrocolony after 36 hours of growth (n=3); (iv) the maximum growth rate monitored during the first 24 hours (n=3); (v) the morphological complexity C-score after silencing of the indicated ncRNA, as shown in figure 6 (n=3). **B**- Statistical data of PCA and HCA (figure 7B). Significant loss of fitness is highlighted.

**Table S3: gRNA_location_list**. **A**- List of targeted genes and transcription units. **B**- sgRNA guide DNA sequences and corresponding genomic location.

**Table S4: PEL I vs NI fitness.** Fitness parameters and their significance for all targets (compared between induced and non-induced conditions) in the pellicle (**A**), with data extracted from significantly depleted (**B**) and enriched targets(**C**).

**Table S5: PEL_I vs PLK_I fitness**. Fitness parameters and significance for all targets in the pellicle compared to the planktonic phase under induced conditions (**A**), with data extracted from significantly depleted (**B**) and enriched targets(**C**).

**Table S6: PLK_I vs NI fitness**. Fitness parameters and their significance for all targets (compared between induced and non-induced conditions) in the planktonic phase under the pellicle (**A**), with data extracted from significantly depleted (**B**) and enriched targets(**C**).

**Table S7: MC_I vs NI fitness.** Fitness parameters and their significance for all targets (compared between induced and non-induced conditions) in the macrocolony (**A**), with data extracted from significantly depleted (**B**) and enriched targets(**C**).

**Table S8 : PEL_NI vs PLK_NI fitness.** Fitness parameters and significance for all targets in the pellicle compared to the planktonic phase under non-induced conditions (**A**), with data extracted from significantly depleted (**B**) and enriched targets(**C**).

**Table S9: Mean depleted targets across biofilm models:** Most depleted or enriched targets: The targets were selected based on a statistically significant fold change (FC) of greater than 2 in either depletion (green, depletion D) or enrichment (red, enrichment U) in at least one of the biofilm models. The models are: Macrocolony (MC), pellicle (PEL) and planktonic phase under the pellicle (PLK). For these three models, the induced condition for dCas9 expression (I) was compared to the non-induced condition (NI). Additionally, the PEL was compared to the PLK phase under the induced condition. Comparing PEL and PLK under non-induced conditions revealed a loss of gene fitness resulting from the system's leakiness under the experimental conditions. For each target, FC>2 values are in bold and, when present in other models, FC>1.5 values are in plain text. These data were retrieved from Tables S4, S5, S6, S7 and S8. Aggregated Q-values are indicated.

**Table S10: Genetic material. A**- list of strains. **B**- Oligonucleotides informations and CUTE-dcas9-GFP module DNA sequence.
